# Maximum likelihood point estimates for improved population genetics statistics

**DOI:** 10.1101/2025.08.01.668180

**Authors:** Andrey Vaulin, Jarkko Salojärvi

## Abstract

Likelihood-based approaches that incorporate the uncertainty in basecalling have become a standard approach in genotyping. Likewise, the accuracy of population genetic estimators could be improved if these uncertainties could be taken into account. Moreover, the classic asymptotically unbiased estimators for population genetic parameters may yield impossible values with finite data, and introduce a median bias driven by sample size, which complicates the comparison of populations of different sizes. Here we develop an approach (https://github.com/anulin/PiThetic) that more accurately estimates the population genetic parameters and addresses the question of comparison (median) bias. Our approach improves by directly estimating population genetic parameters from basecall accuracies in terms of Phred scores, instead of using allele frequencies. We provide three improvements to classic statistical estimates: (i) we apply Phred scores to improve the accuracy of the estimates, (ii) we show how the pursuit for an unbiased estimator can lead to impossible and/or less accurate values, and provide a corrected estimator, and (iii) we provide a new estimator that minimizes the median bias for nucleotide diversity and another estimator that reduces error for D, because the standard Watterson θ and Tajima’s D estimators have high error. Overall, this work facilitates the more accurate estimation of population genetics statistics from sequencing data.

## Introduction

Standard variant calling uses maximum likelihood (ML) approaches to call genotypes from aligned read data (for example ANGSD ^1^, GATK ^2^ and FreeBayes^3^), and these are used in turn for estimating population descriptive statistics such as allele frequencies and population genetic statistics with tools such as pixy ^4^, PLINK ^5^, or vcftools^6^. However, such approach causes bias and possible error^7^ as the uncertainty in the genotype calling is not carried through to the estimated statistic. In shotgun sequencing, the probability of an erroneous nucleotide call at a certain a position in a read is provided as a Phred score^8^. Knowing the uncertainty of basecalling can help improve accuracy of subsequent estimates, and some works have already tried to incorporate base quality scores directly into population genetic statistics estimation^1,9,10^. However, the proposed methods are often oversimplified or applicable to some limited cases. Also, while statistics obtained through basecalls are often more important than mere counts, to our knowledge there are no tools that calculate population genetic statistics, other than site frequency spectrum^1^, directly from Phred scores. Here, we propose a method that utilizes the Phred scores to calculate more accurate statistics and, although the focus here is on the biallelic case, it can be extended to multiallelic case as well.

Different population genetic statistics are used to draw conclusions about the evolution of the population. For example, nucleotide diversity (π)^11^ is linked with the effective population size (N_e_), and it can be also used to detect selection by comparing the diversity in synonymous vs non-synonymous sites (π_n_ vs π_s_ ratio)^12^. However, estimating the diversity from a population sample is problematic. With finite data, due to the correction for the degrees of freedom lost in its estimation, the standard asymptotically unbiased estimator provides values where the range of the estimates depends on the sample size, with larger range for a smaller sample. Additionally, small samples drawn from the same population have a high variance, that is, a higher probability of producing more extreme values. The estimation may also result in impossible estimates, because a relatively small sample is used to make conclusions about the whole population, and smaller sample sizes have a larger positive bias. Such bias would have an effect on statistical significance (depending on the test) for smaller samples, which may affect the results. An illustrative example would be the case when the true π is close to 0.5, as here the standard estimator yields impossible values which makes the estimator inappropriate. For instance, a sample of 12 haploid individuals (or 6 diploid), may reach π values of 0.54, while 50 haploid individuals can reach 0.51, both of which are impossible values, as the theoretical maximum for π is 0.5. Such cases are obtained because the estimate 2*p* − 2*p*^2^ is multiplied by a correction term 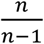 to make it unbiased. Here we propose an MLE method for nucleotide diversity that is not affected by these issues. We also directly implement Phred scores into the calculations instead of having to estimate frequencies as an intermediate step to improve accuracy.

Another population genetic statistic is Watterson θ. It is used to make conclusions about mutation rate and effective population size, and it is used in tests for selection such as Tajima’s D. However, most values from the standard estimator are below the true value, the estimator has a very large variance and decreases very slowly with sample size, altogether making it impossible to get an arbitrarily small error by increasing sample size. Some methods have been proposed to improve the accuracy^13–15^ but they all rely on site frequency spectrum. Site frequency spectrum is affected by selection and therefore such improvements make these estimators not useful, especially when testing for selection. Tajima’s D in turn depends on θ and mutation rate which causes error in estimating selection. We discuss the MLE for θ and an alternative selection test for Tajima’s D.

## Results and discussion

### Nucleotide diversity π and its traditional estimation

Nucleotide diversity π_*d*_ is defined by ^11^:

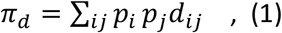

where *p*_*i*_, *p*_*j*_ are the frequencies of sampled allelic variants *i* and *j* in the population, and *d*_*ij*_ is the per nucleotide difference between alleles *i* and *j*. An unbiased estimator of nucleotide diversity is obtained with the classic formula ^11^:

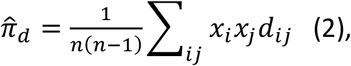

where *x*_*i*_ and *x*_*j*_ are the counts of alleles *i* and *j, n* is the sample size – number of haplotypes in the sample. Normalization by 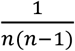 results from a degree of freedom bias correction for the frequency-based estimator of π that can be derived from (1), and is essentially the number of possible pairs of individuals. However, with small sample sizes this correction causes asymmetric distribution around the actual value of the original population, biased towards greater values. With small sample sizes the estimators have different ranges depending on the size of *n* and, hence, incomparable results when the sample sizes differ. For example, at the extremes, for a population size of 2 the value ranges from 0 to 1 while for an infinite population the value ranges between 0 and 0.5. This hampers the comparison of estimates derived from populations of different sizes.

Being able to compare values for different sample sizes and experiments is of high importance for any score and estimator, and therefore we next develop an estimator in which the value range and error type are independent of the sample size.

### ML-based nucleotide diversity estimator

In matrix form, formula (1) can be written as

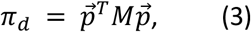

Where

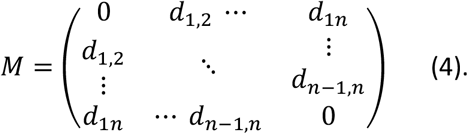

Thus, for a given value of π, the frequencies make a hypersurface of second order (**Figure 1**):

**Figure 1.**
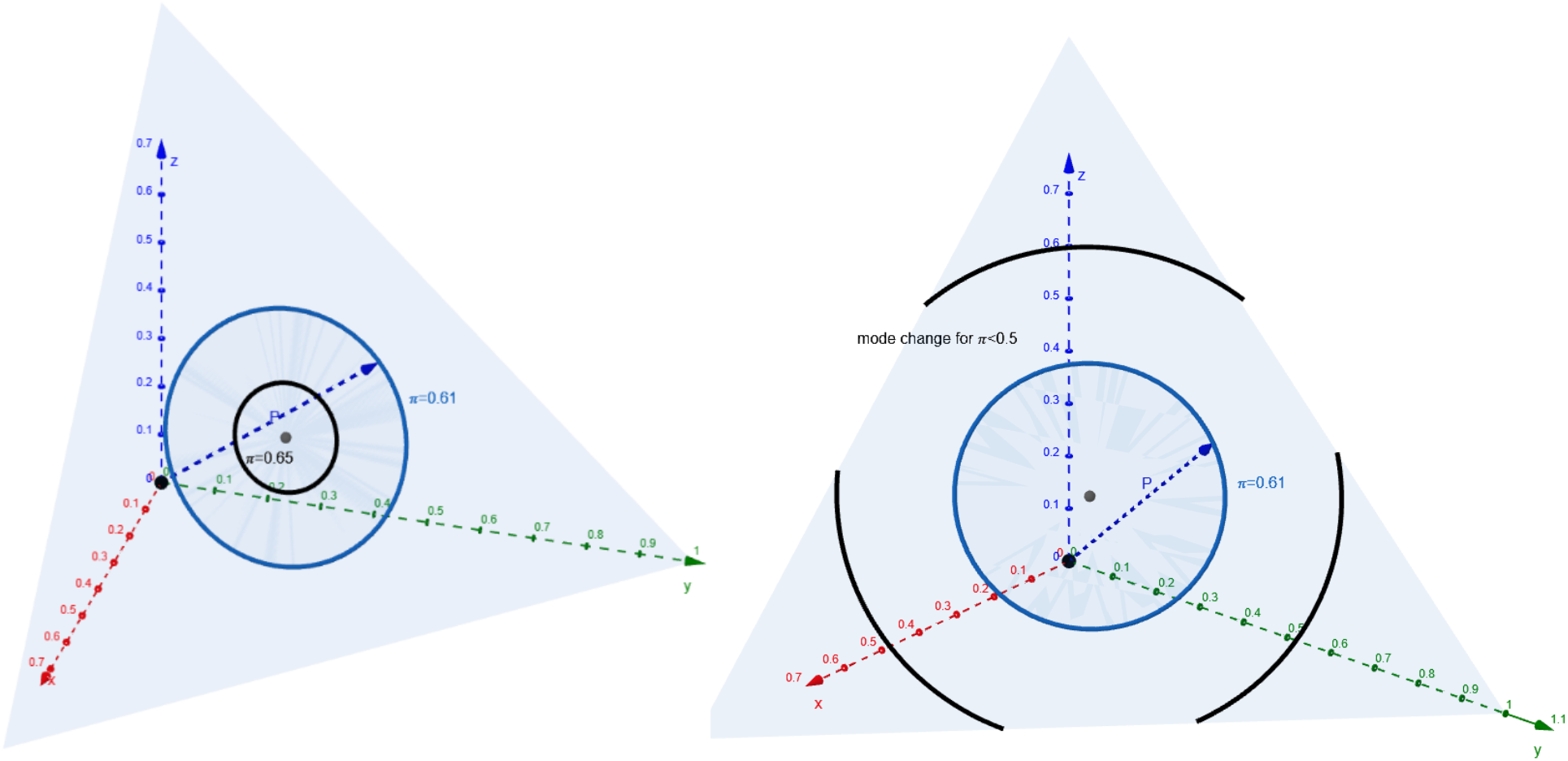
Three-dimensional example of *S*_π_ (circles) for different π. Axes correspond to allele frequencies. Triangle surface is the possible sets of frequencies (their sum equals 1). Left: figure for a larger value of π. Right: *figure* with mode change for smaller π. The mode change is due to boundary conditions for frequencies. Circle length (parameter space) no longer grows linearly with π going down. There will be n-2 mode changes for n-allelic site.

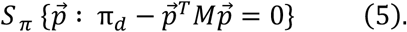

Since a given π_*d*_ does not provide the full information about 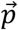 and only limits possible values of *S* _π_, likelihood for π_*d*_ is an average over 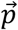 that produces π_*d*_ within the hypersurface:

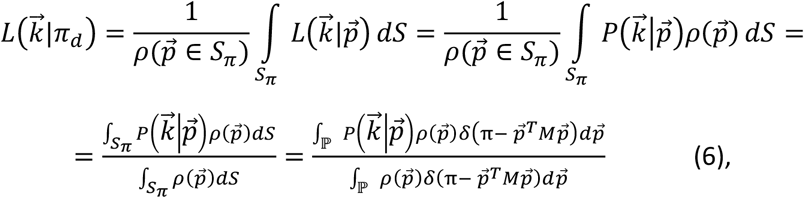

Where 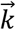 is the vector of counts (the observed data), ℙ is a space of possible values for 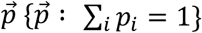 and ρ is the probability density of 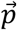. So, to get a maximum likelihood estimate 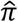 we need to find the maximum average likelihood for 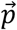

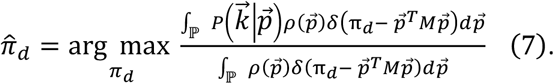

### Biallelic case

Assume the counts to follow a multinomial distribution and the parameters to follow uniform *a priori* distribution (that is, 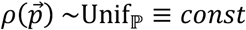) in order to calculate 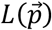.

For the case of a single biallelic site (i.e. *d*_*ij*_ = 1 for i ≠ *j*), the allele probability vector reduces into 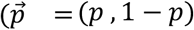, and thus Eq. (3) to π = 2*p* − 2*p*^2^. Therefore the solution to *S* _π_ are roots of the equation π − 2*p* + 2*p*^2^ = 0, i.e., one or two values, and applying (6) we get the corresponding likelihood in case of observed counts m, k:

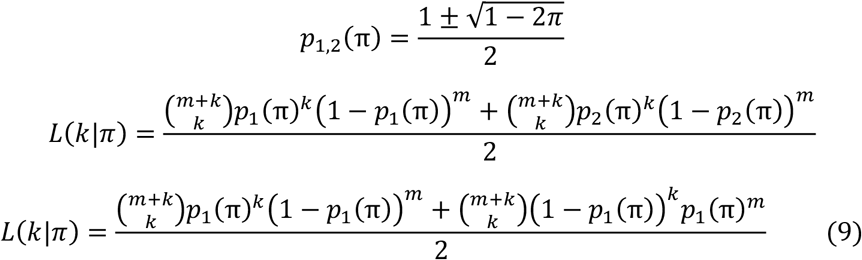

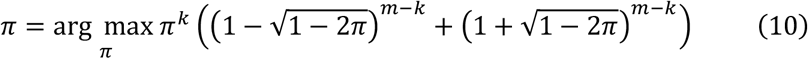

The estimator median follows the true value (i.e., is median-unbiased) and therefore does not have probability dependence on the sample size.

### Simulations for ML-based estimator of π

For an empirical validation of the MLE-based π, we simulated data assuming an infinite population (i.e., sampling with replacement) and binomial/multinomial (four-allele) allele frequency distribution. For a sample size of 40 individuals, the possible frequencies *p* arising from allele counts range between [0,20] (altogether 21 different values of p). However, there are only 18 possible values for the MLE π in the case of 40 samples, because several different counts produce the same value of MLE. Samples of size 40 were generated 50,000 times for the 18 different binomial distribution parameter values *p* corresponding to these values of π, followed by calculation of MLE-based and the standard π estimators.

The box plots of the estimates were plotted for all values of *p* in the grid to visualise the errors and deviation of standard scores (estimators) median and then compared with the proposed ways of improvement (**Fig. 2, Fig. 3**).

**Fig. 2.**
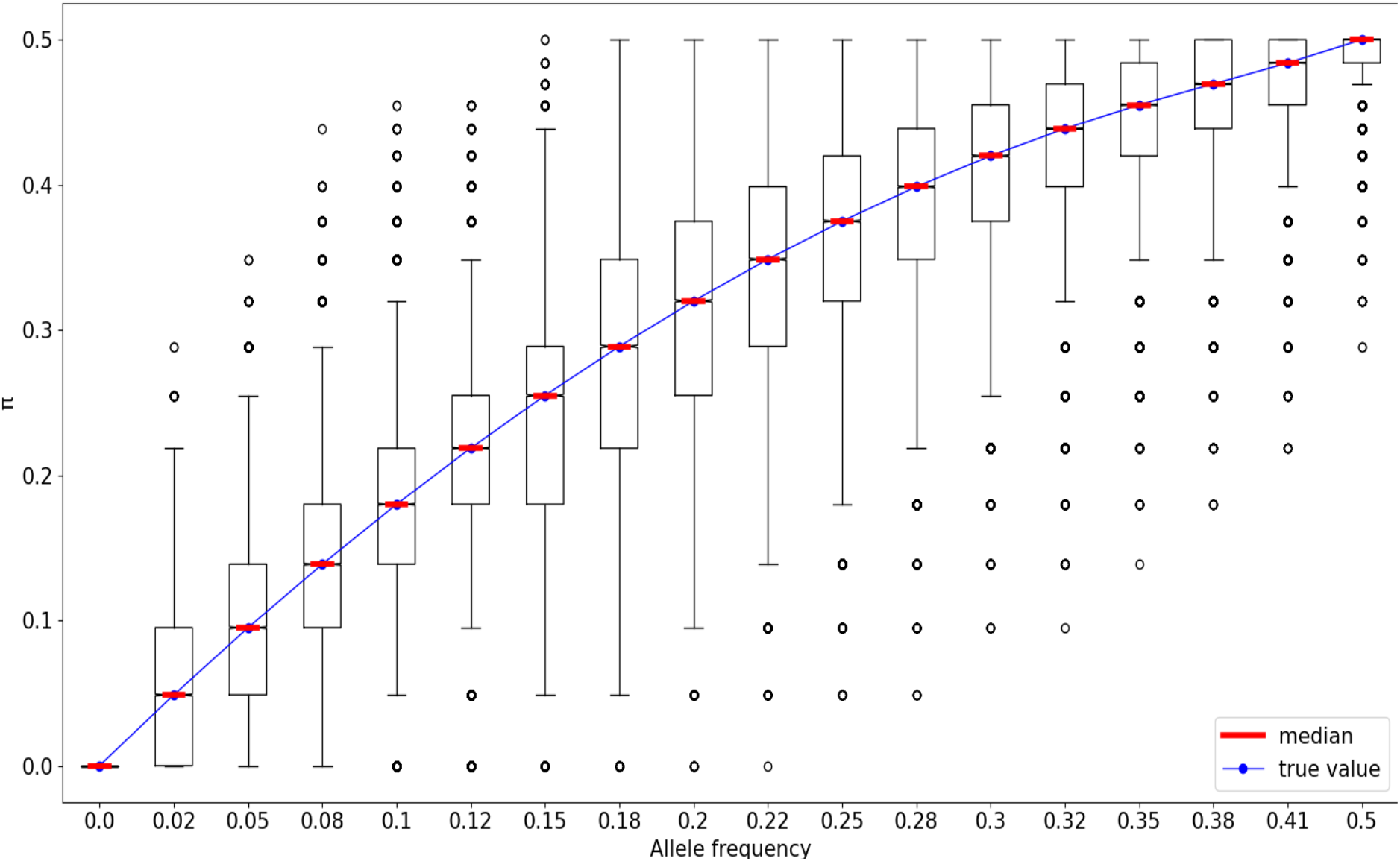
Boxplot of MLE π values for different frequencies for sample size of 40. Blue dots are the true values of π.

**Fig. 3.**
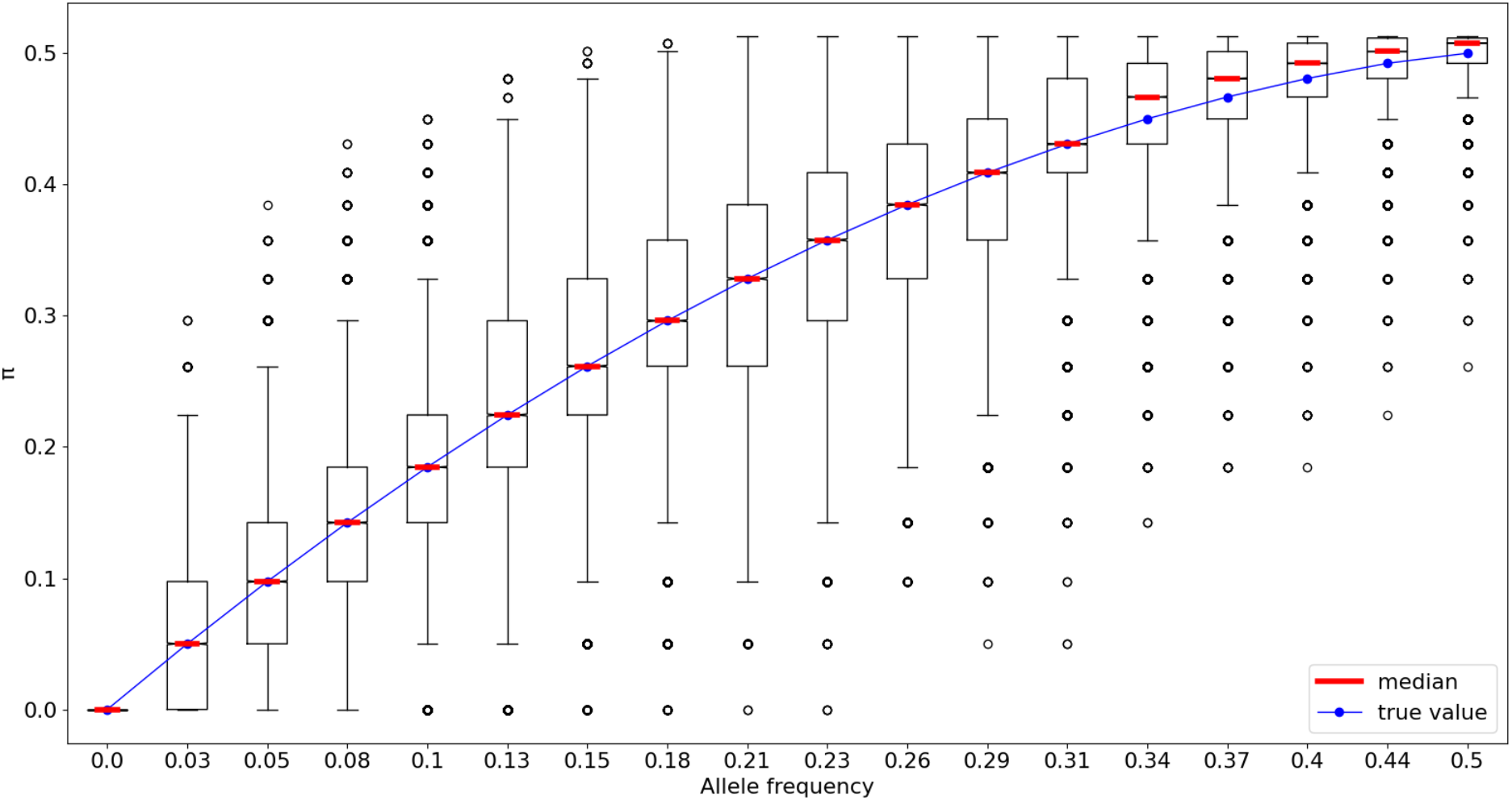
Boxplot of bias corrected π values for different frequencies for sample size of 40. Blue dots are the true values of π.

The simulations display that MLE median clearly follows the true value and for the standard estimator median becomes higher than the true value after frequency ≈0.33. Also, impossible values of π>0.5 are observed for the standard estimator, though π ≤ 0.5 for a biallelic site. In contrast, for our MLE-based π the median follows the true value.

### Phred score-derived MLE for allele frequency and π for diploid organisms

We next extend the MLE-based estimator to include ploidy and basecall accuracies in terms of Phred scores. Assume the confidence of a correct call to be *h*_*i*_ ∈ [0,1]. Then the binomial distribution of allelic nucleotides in individuals together with ***h*** defines the distribution of nucleotides in reads coming from their paired, diploid chromosomes^16^.

For a diploid organism, likelihood for four alleles (*a*=1,2,3,4) and actual frequencies *p*_*a*_, a SNP in each locus in one individual has two possible states: the site is either i) homozygous or ii) heterozygous. Frequency likelihoods for an individual j for these states are

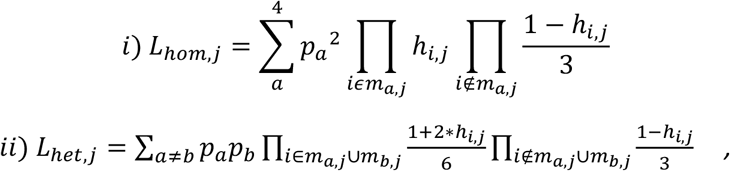

where *m*_*a,j*_ are the indices of reads with certain nucleotide *a* for individual *j*. Because the states are mutually exclusive, the total likelihood for an individual *j* is their sum and the total likelihood for the sample is thus a product of likelihoods for different individuals:

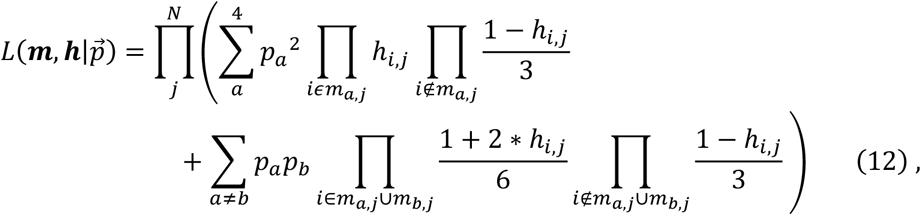

where *N* is the number of individuals. For biallelic frequency there are just two alleles *a* and *b*, and their frequencies are *p*_*b*_ = 1 − *p*_*a*_:

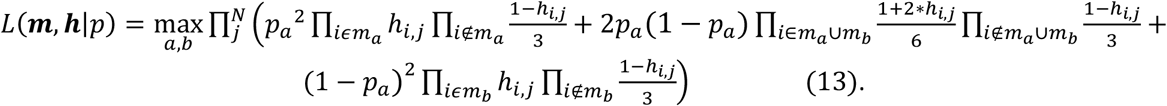

For a biallelic site, π follows the same logic from (9):

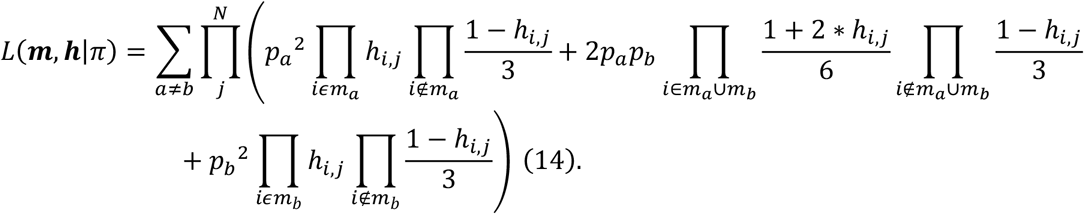

### Simulations for phred scores and diploid data

To simulate data, we sampled a set of 60 Phred scores from real-life sequencing data (distribution shown in **Supplementary Fig S1**), such that there were lower scores present in the sample. SNPs were assumed to be biallelic. Random values from binomial distribution with a probability *p* were sampled to get true sample allele frequencies. Phred scores were sampled randomly without replacement, and the erroneous calls were sampled from a Poisson-Binomial distribution, with the probabilities defined with the sampled Phred scores as parameters, and applied to each individual to simulate the observed reads. Errors are a particular problem for smaller datasets with lower coverages, therefore diploid samples from a population of 15 individuals with sequencing coverage of four were generated. Data generation was done 100,000 times for ten different values of *p* in an interval [0,0.5] with increment of 0.0(5). Likelihoods were calculated for the data using the Phred scores. Medians of standard (SNP call) allele frequency, MLE for frequency (13) were compared and ML frequency estimates are more accurate than the standard estimates. Similar comparison was done for π from formula (14), π calculated from MLE frequency and the standard π. Again ML π estimates (14) are the most accurate. Simulations for random Poisson-distributed coverage (λ=4) were also made, with similar results; see (**Supplementary Fig S2-6)**.

A comparison of the medians of observed frequency, MLE for frequency, and the actual value (**Fig. 4**) shows that MLE frequency is better in terms of median compared to the naïve estimator.

**Fig. 4.**
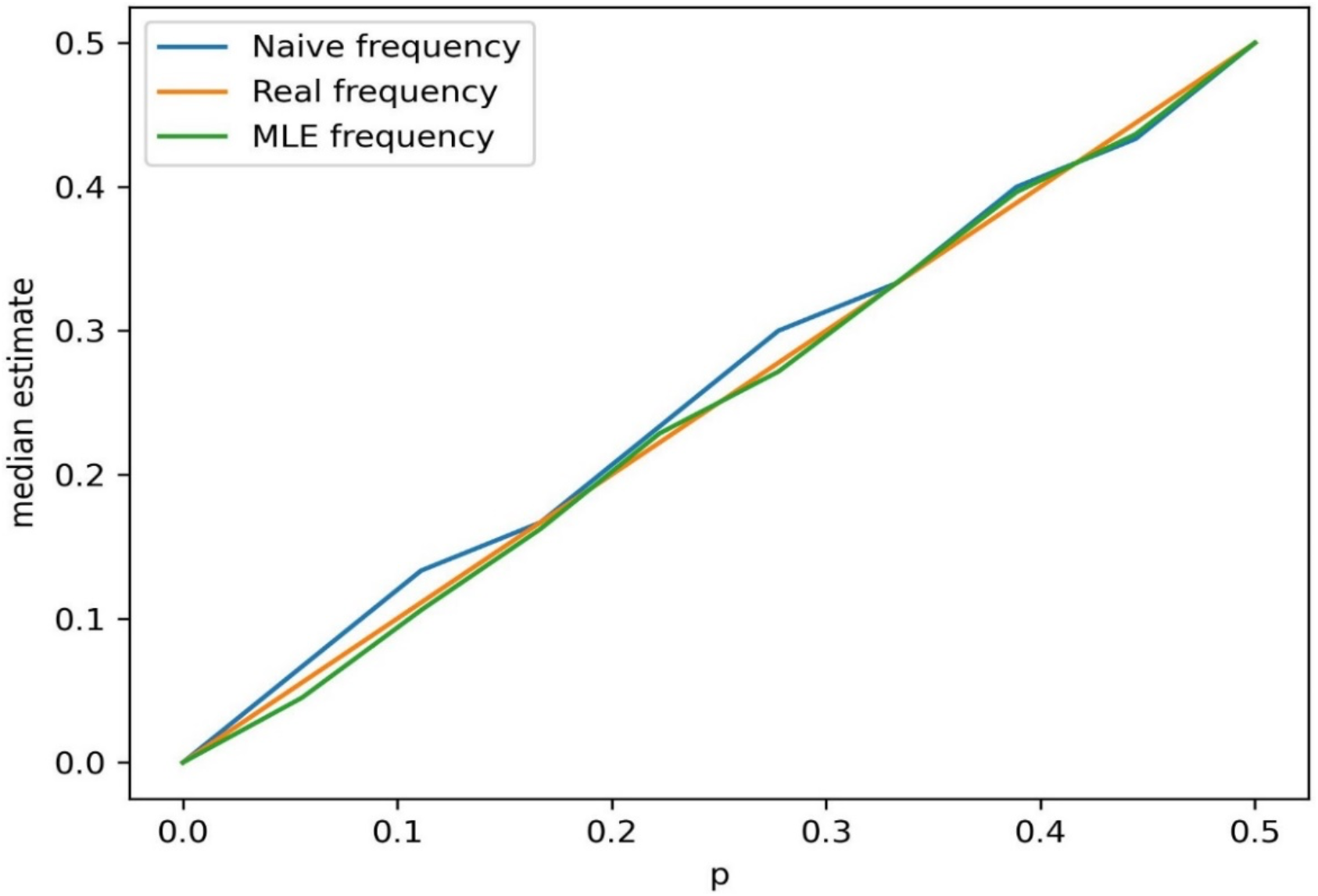
MLE for frequency derived using phred scores. The MLE frequency is better than the standard one.

A visualization for the mean error to the true π for the different estimates difference (**Fig. 5**) showed that π MLE calculated directly from Phred scores is the most accurate compared to the naïve π and π calculated from MLE frequency.

**Fig. 5.**
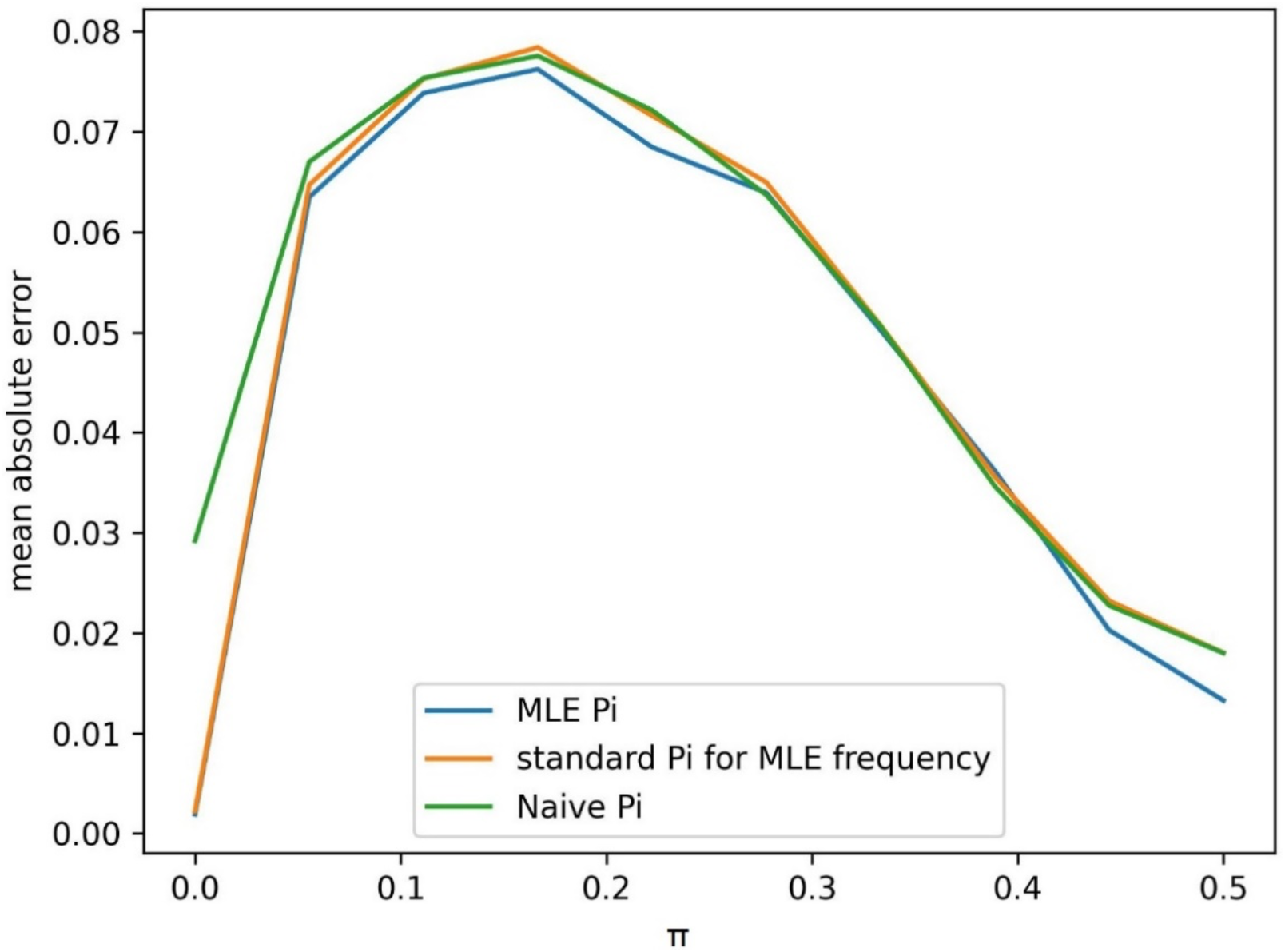
Mean absolute errors for MLE for π derived using phred scores(blue), for standard π calculated using MLE frequency (orange) and for naive estimator based on observed frequency (green). MLE π has the lowest error among the estimators

### Multi-nucleotide π

Nucleotide diversity π of a region of size *l* is the average of nucleotide diversities π_1,2,…*l*_ of nucleotides within that region. Neither sum of MLEs nor sum of median-unbiased values does necessarily produce MLE or median-unbiased estimator of the sum. Instead, we must calculate them separately. Likelihood of π is then:

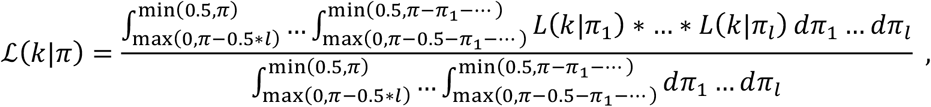

where the integral range from 0.5 results from biallelic π. For four alleles the value should be changed to 0.75. For larger regions, especially with many SNPs, a straightforward calculation of this integral becomes computationally hard. However, for these regions (approximately when region size>sample size) the standard estimator becomes median-unbiased (given there are no Phred-score or other errors) and is easier to compute, given a known sample frequency. Faster computation results because of the general properties of an unbiased estimator, meaning that simple averaging produces an improved estimate. For the case when there is uncertainty in data and fast computations are needed for regions, we provide another estimator below.

When sample allele frequencies are not known or are known with different accuracy (e.g. base call and ploidy-associated errors), it becomes important to normalize population frequency estimates by estimator variances before taking the average, otherwise the accuracy may not increase with the growing region/sample. When the organisms are diploid there can be three possibilities: individual is homozygous for reference allele, for alternative allele or heterozygous. The respective probabilities for a given allele pair *ab* are:

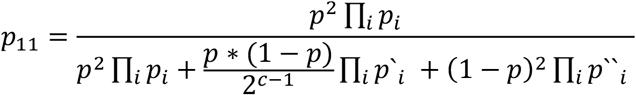

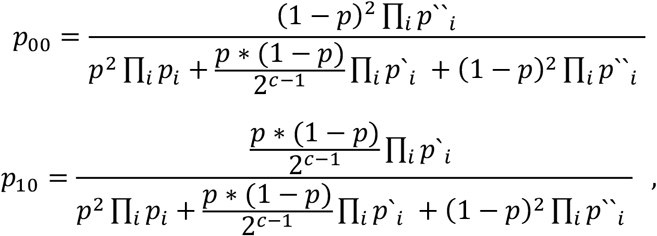

where *p* is the sample allele frequency, *p*_*i*_ is the probability of observed read given reference allele, *p*‘_*i*_ is the probability of observed read given reference or alternative allele and *p*‘‘_*i*_ is the probability of observed read given alternative allele. The expected number of alleles in an individual is 2p and

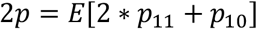

Therefore, the allele frequency can be estimated as the root of the following equation:

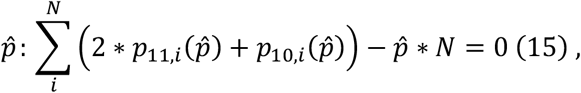

where N is the number of individuals. The nucleotide diversity is now

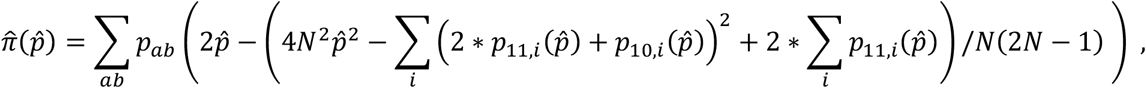

where *p*_*ab*_ is the probability of an *ab* allele pair:

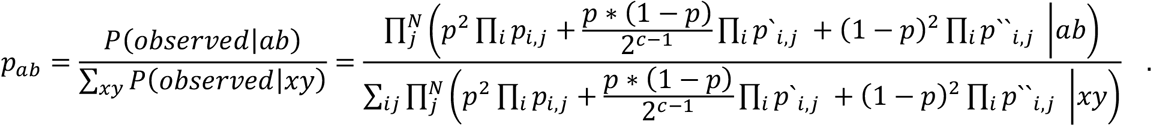

Variance is then estimated as:

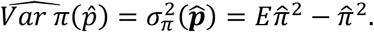

Expectancy here is taken over possible basecall errors (conditioned on 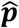, which is a vector of frequency estimates for different allele pairs). Now, a better estimator of 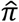 in a region is produced by normalizing for this variance:

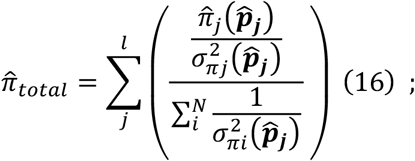

here *j* is position in the region.

### Maximum Likelihood estimation for θ

θ is a parameter of coalescent process and it describes the product of the effective population size and the mutation rate. Because in neutral coalescent process the expected sum of nucleotide diversities is equal to θ, this sum (also called π) is sometimes used as an estimator of θ, effective population size N_e_, or mutation rate μ, though the estimator is not consistent - even with large sample sizes, π does not converge and has above 0 variance. Watterson estimator, however, does converge.

We follow the classic Watterson’s derivation^17^ for θ and consider the Wright-Fisher model of evolution (**Fig. 6**).

**Fig. 6.**
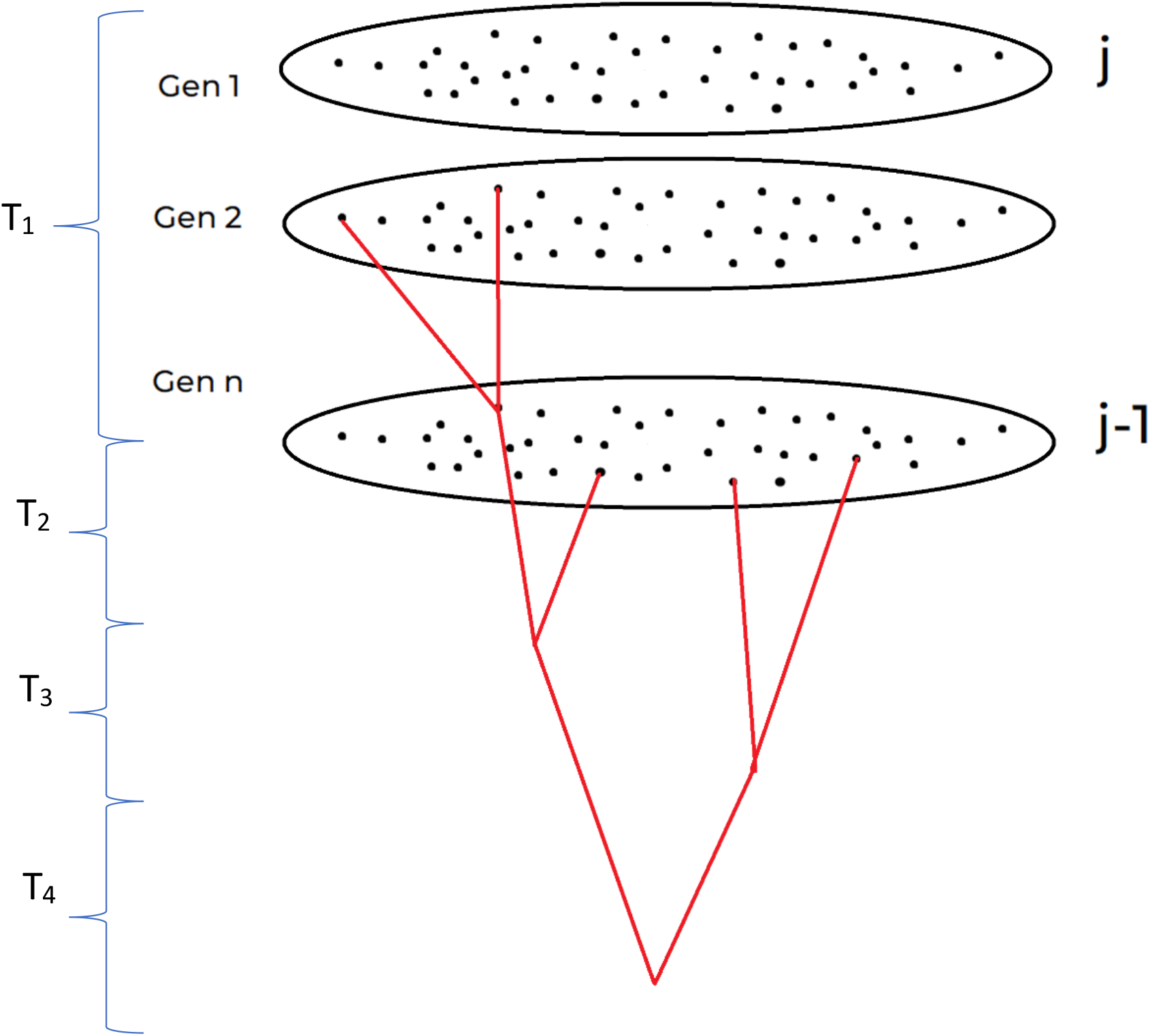
Scheme of Wright-Fisher model.

Given a population of constant size 2N_e_ the probability of coalescence for individuals *i* from (and assuming i<<2N_e_) is

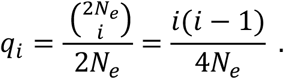

The time until coalescence *T*_*i*_ is then a geometrically distributed variable *X*_*i*_ with parameter *q*_*i*_.^17^ Next, assume a sample of *j* haploid individuals from the population with 2N >> j.

The mutations in one lineage in a generation are assumed to have Poisson distribution with parameter λ=μ, the mutation rate. From the properties of Poisson distribution it follows that the sum of Poisson distributions with means λ_1_, λ_2_, λ_3_ is a Poisson distribution with mean λ = λ_1_+ λ_2_+ λ_3_. Then, total number of mutations between *i* individuals during the time *T*_*i*_ follows Poisson distribution,

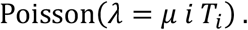

Because *T*_*i*_is not known and has geometric distribution, the number of mutations *K*_*i*_ between i individuals during *T*_*i*_ has compound geometric-Poisson distribution

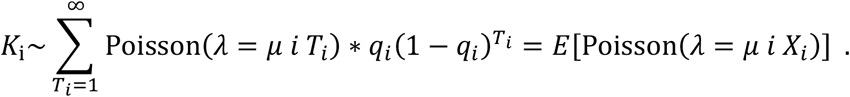

Therefore the total number of mutations *K* between the *j* individuals in our sample is distributed as sum of compound geometric-Poisson values:

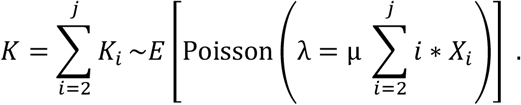

Therefore, given the observed number of mutations *k*, an MLE can be obtained by taking the derivative of the expectation and setting it to zero:

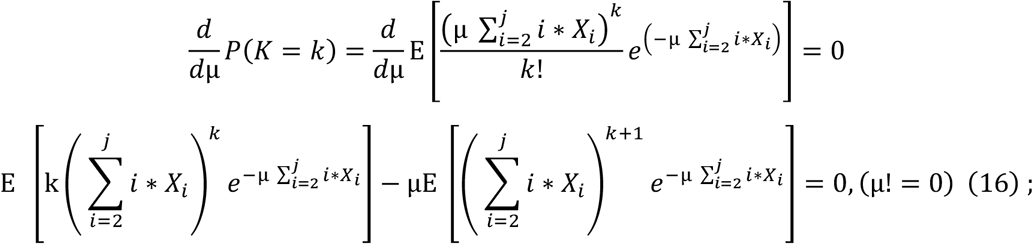

The *k*th moment of geometric distribution is 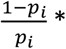 Polylogarithm_-k_(***p***_***i***_) =

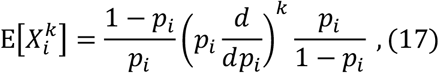

And

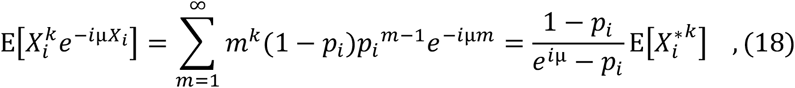

Where 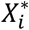 is geometrically distributed *p*_*i*_^∗^ = *p*_*i*_*e*^−*i*μ^. Now, using (16) and (18) we get

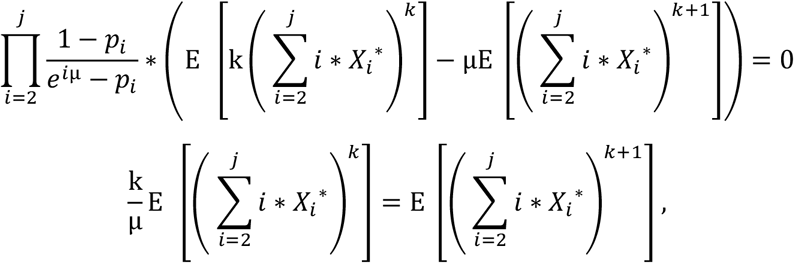

and then applying (17) yields

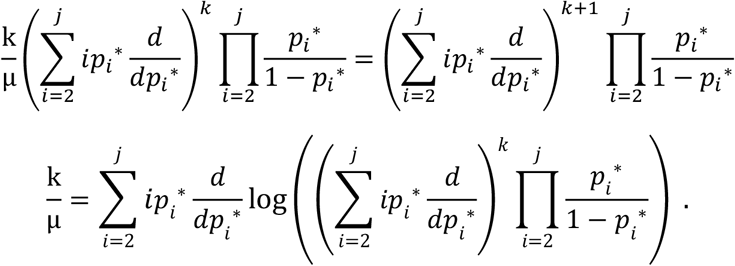

Since 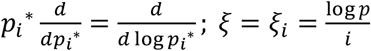, then:

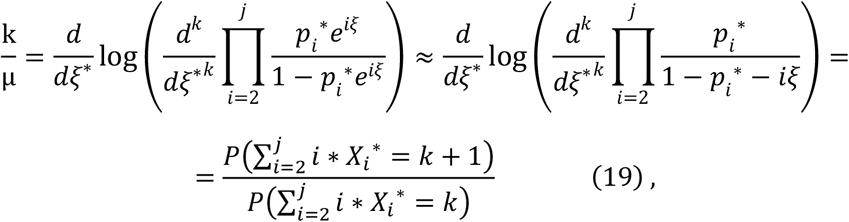

which provides us mutation rate estimate μ.

Then, given N_e_, we can get *θ* with standard formula

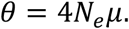

For large *k* and *j* the computation may take very long, and for this case we propose to solve the expectation (14) numerically, e.g., by doing simulations.

### Median-unbiased estimator

However, the MLE estimator suggested above is median-biased. For median-unbiased estimator we need cumulative distribution function to be ½:

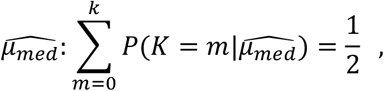

by the definition of median. Then we can instead get the parameter μ as a solution to the following equation:

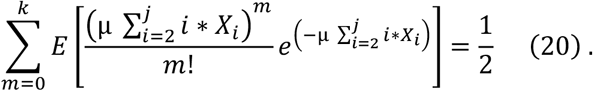

Wright-Fisher model relies on the assumption that there is no recombination. In case of strong recombination (0 linkage disequilibrium) each mutation has its own ancestral tree, and the number of possible mutations equals the number of loci. The distribution would then be binomial with the size parameter *n*=number of loci, but since the number of loci is assumed to be infinite, the distribution is Poisson, 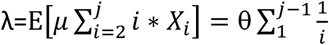. In this case Watterson estimator is the same as the MLE. Median-unbiased estimator is a solution to equation:

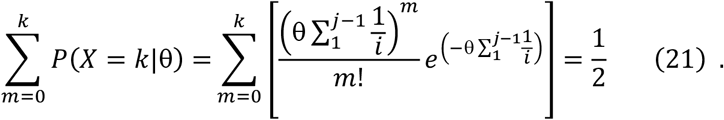

### Simulations for Watterson θ and Tajima’s D

The mutation rate *μ* was assumed 10^−5^ mutations per generation and population size *N*=10^6^. The times to coalescence *T*_*i*_ when exactly *i* lineages existed were generated from geometric distribution with parameters 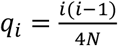, and the number of mutations were drawn from Poisson distribution with 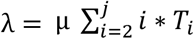. This was done 200,000 times and then the medians were calculated for Watterson θ and MLE Watterson θ. The results illustrate that accuracy of estimators increases very slowly after sample size of ∼20. Error is 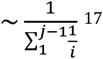.

While standard estimator tends to produce values that are below the true value, MLE generally produces greater values (**Fig. 7**). Perhaps a less known property of the Watterson estimator is that the standard estimator converges to the true value very slowly, with the error being 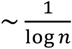 where ***n*** is the sample size.

**Fig. 7.**
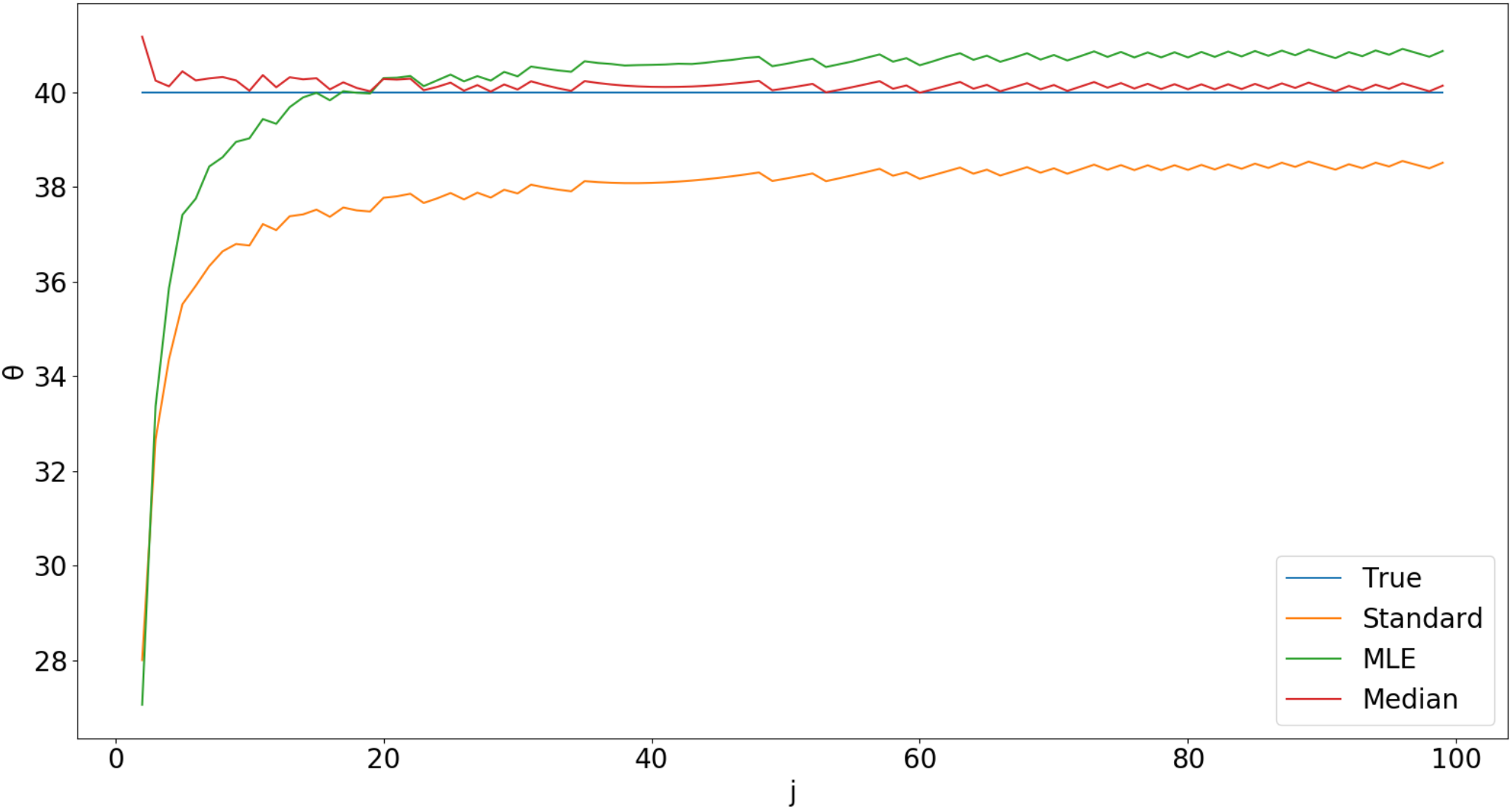
*MLE (green), standard Watterson* θ (orange) and median-unbiased estimator medians. The true θ of 40 is the blue line. Estimators converge to the true value very slowly.

This requires unrealistic sample sizes (in the order of 10^10^) to reduce the error to 1% or less. A real-life scenario with a sample size of 100 produces 5% downwards median bias (i.e. in half of cases the estimator is more than 5% lower than the true value).

However, the MLE method reaches the plateau of approximately accurate values faster than standard average-based estimator, already with sample size of 10 (while standard requires 20) and the trend continues (at least up to sample size of 1000) with notably smaller difference between median and the true value. The variance of the estimator does not change much, but is slightly higher for the ML estimator (**Fig. 8**).

**Fig. 8.**
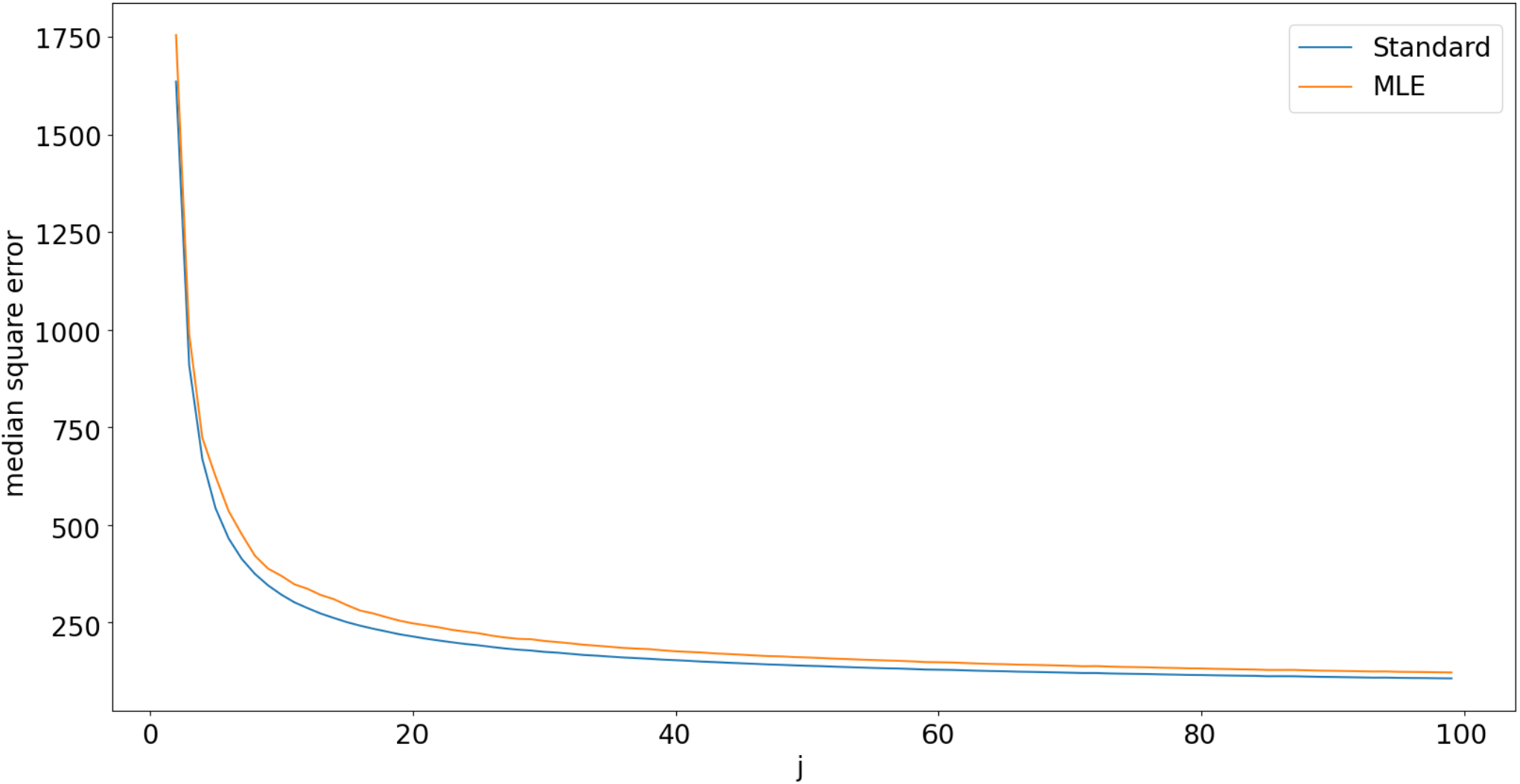
MLE Watterson variance vs standard estimator variance. ML estimator variance (orange) is generally slightly higher than standard estimator (blue)

There exists previous work on improving Watterson θ ^13,14^, and we next compared those to our MLE. The Futschik’s unfolded BLE estimator clearly has greater median bias, with smaller median values than both MLE and standard estimator (**Fig. 9**).

**Fig. 9.**
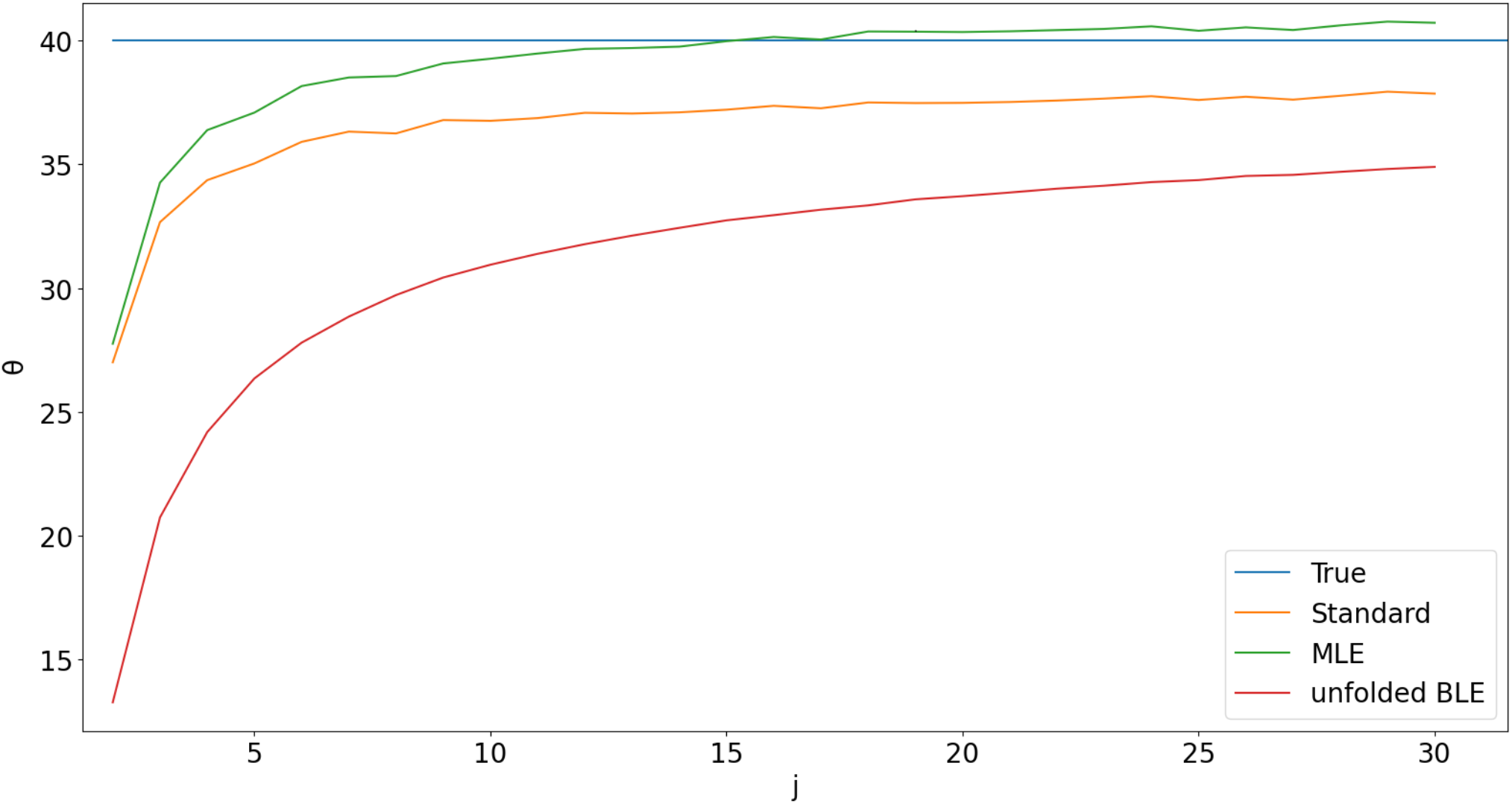
Futschik’s best linear estimator(BLE) compared to others. It has the largest median bias

Furthermore, despite having smaller mean square error, it has larger median square error than both MLE and standard estimator in case of strong recombination (**Fig. 10**) and for smaller sample sizes when there is no recombination (**Fig. 11**). This is because square error penalizes for the largest errors, while paying less attention to smaller errors or majority. In contrast, the median error pays attention to a majority of results while ignoring separate cases/outliers.

**Fig. 10.**
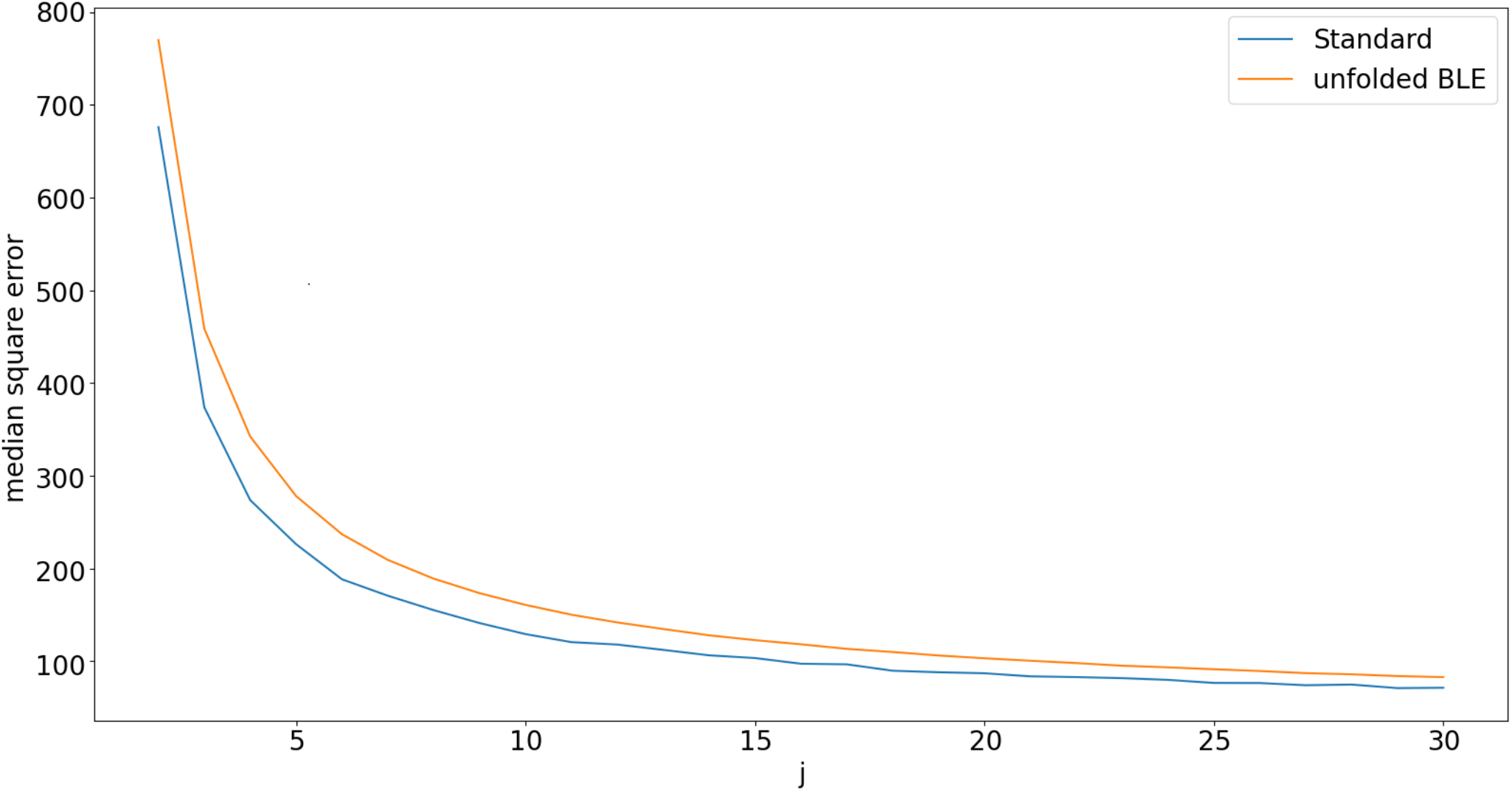
Median square error for BLE and standard estimator (with recombination)

**Fig. 11.**
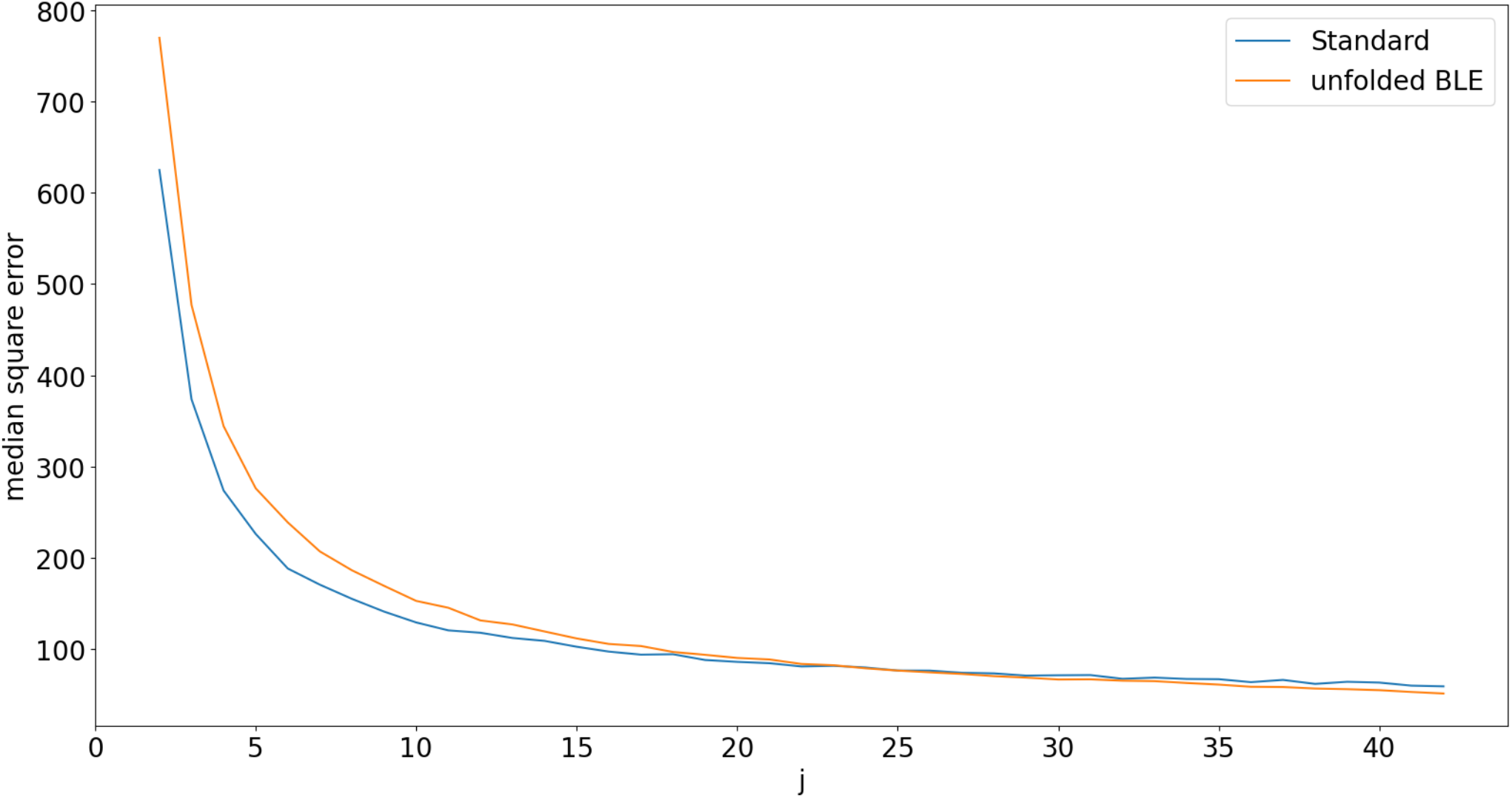
Median square error for BLE and standard estimator (without recombination)

Fu’s BLUE estimator behaves the same way as standard in terms of median and median error if there is strong recombination ^14^. Both Futschik’s and Fu’s estimators rely on the assumption that there is no recombination and do not provide any improvement for the median error if the recombination is strong and each mutation has its own phylogenetic tree.

Finally, the MLE median error is generally slightly smaller than the standard estimator’s median error (**Fig. 12**). Therefore BLE is the best estimator in terms of median square error for θ when sample size is >∼33 if there is 0 recombination, while MLE is best for smaller sample sizes.

**Fig. 12.**
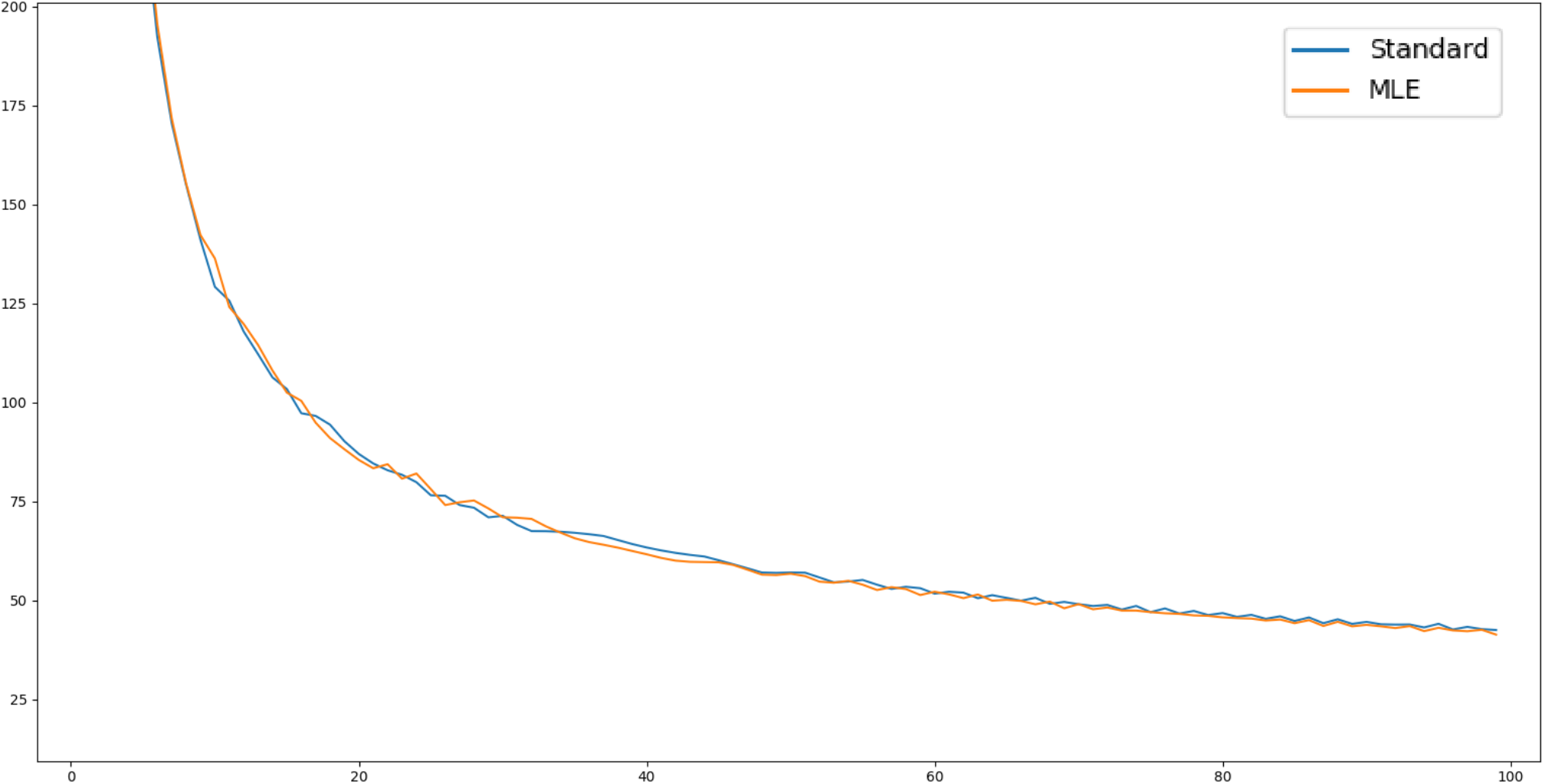
MLE and standard estimator median square errors are very similar.

### Tajima’s D

Tajima’s D is calculated with the following formula:

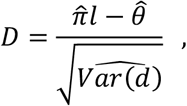

where 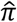 is standard nucleotide diversity estimator calculated within a region, *l* is the region size, 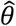 the Watterson θ estimator for the same region, and 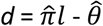, if 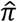 is derived from coalescent theory (i.e. 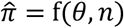) ^18^.

Under neutral model without recombination and constant population size, median of *d* is below 0, and thus the median of Tajima’s D is negative because 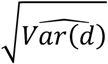 is always positive. Moreover, 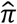 is not consistent^19^ (does not converge to θ), which means that the negative error cannot be fully removed. However, when the recombination is strong the median of *d* is 0. Watterson 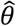 is a function of the number of mutated sites which does not strongly depend on selection and adds noise to the statistic due to unknown mutation rate. On the other hand, site frequency spectrum does not depend on the (unknown) mutation rate but strongly depends on selection. Under neutral model with no selection and constant population size the expected site frequency spectrum is known and thus yields an expected per site nucleotide diversity π_*e*_, which under Wright-Fisher model will be 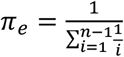. This means that populations with the same history and selection and different mutation rate may have different results for Tajima’s test. To reduce noise coming from the random mutation number *K* and to solve this issue, we propose to remove the dependence on the number of mutations from the formula and treat *K* as a known value. Then the improved test is

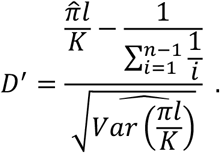

Here 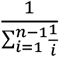 is a known value and has no variance unlike 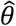.

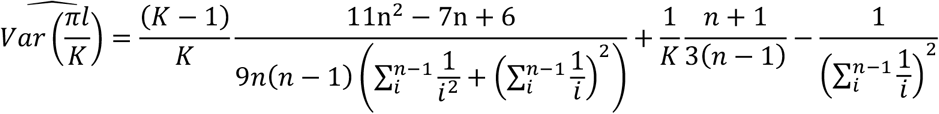

This test is more sensitive and can detect weaker selection signals (or population size changes). Furthermore, it is median-unbiased (**Fig. 13**).

**Fig. 13.**
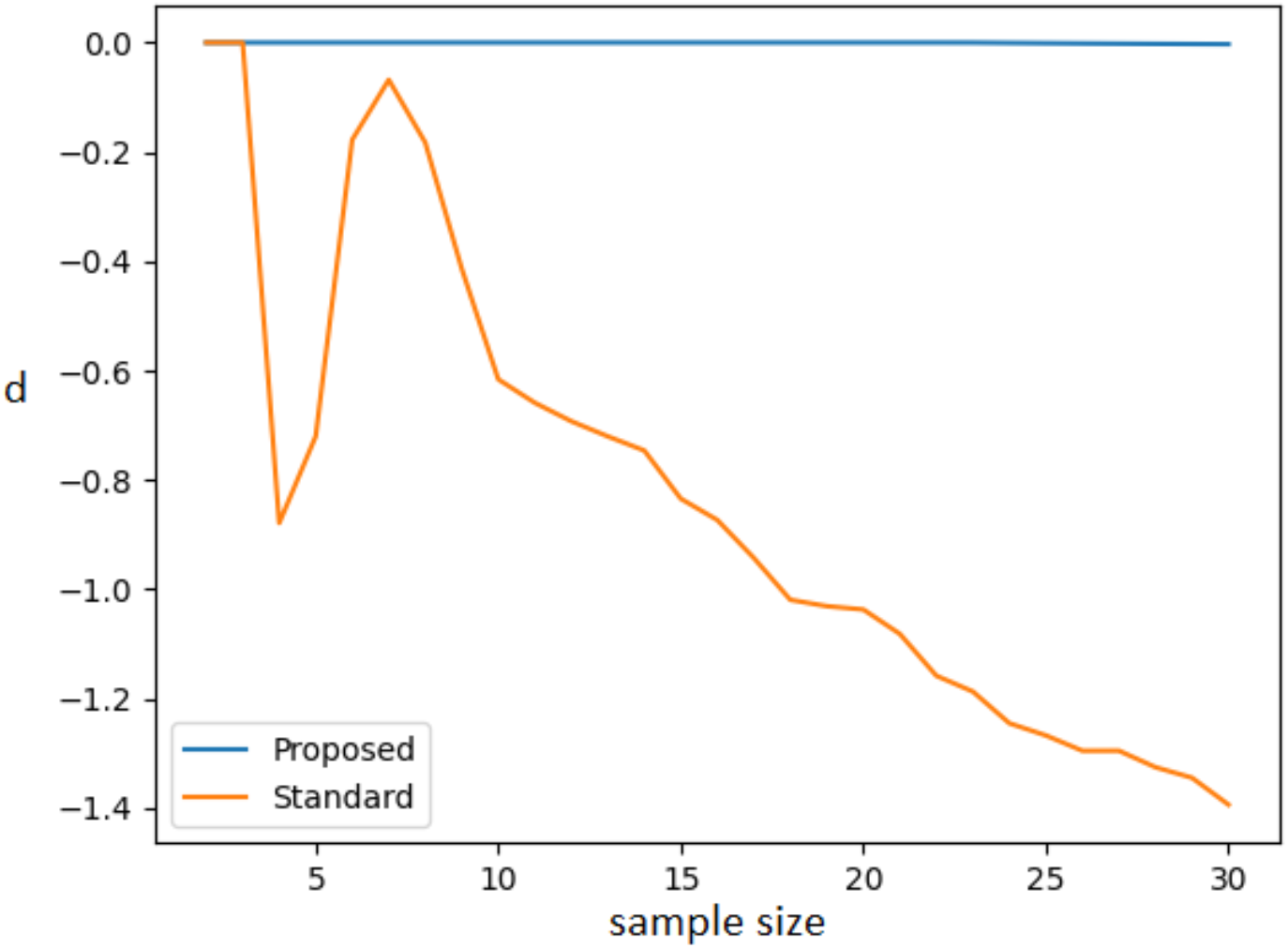
Median of d, the numerator of Tajima’s D estimator. The data was simulated from a population with constant size and no recombination and therefore the correct value is 0. For each sample size (x-axis), 200,000 simulated data sets were sampled and the value of d was calculated. The blue line shows the median value of the new estimate proposed here, the orange line illustrates the median value of the standard estimator.

## Discussion

Here we presented a new set of maximum likelihood-based estimators for population statistics. The proposed methods were shown to be more accurate than the standard estimators. With finite samples, the standard estimators were shown to produce impossible values and were shown to depend on the sample size. Because representative samples are used to study the overall population behavior, this dependence cannot be allowed and, furthermore, it can cause impossible values for the estimated statistic. Moreover, standard estimator’s median can depend on the sample size for the samples drawn from the same population. For standard estimator of nucleotide diversity, smaller sample size increases the median, meaning that most values will be larger than the median of a larger sample and the true value, causing smaller samples to overestimate the true value and thus produce greater diversity estimates than larger samples, even if they would be drawn from the same underlying population. These issues are caused by the pursuit for asymptotic unbiasedness – an approach which requires an average of estimators for the same sample size to be a more accurate estimator. However, it is only useful to simplify the calculations and, given the issues it produces, we believe that it should be dropped. Statistics and scores in most cases in science are used for comparison to make conclusions (for example using *p*-values) and if we are unable to compare them the values have no meaning to us.

Here we suggested an MLE estimator that does not suffer from these biases. For practical purposes, it makes more sense to use median-unbiased estimators. Median is a value such that half of the values in the sample are larger, and half are smaller, and with that it provides us with the information for comparisons of the whole sample. For this reason, we believe that median is more useful and we thus suggest using median-based methods instead of average-based that are asymptotically unbiased. It is important to point out that unlike mean-unbiased estimator, sums of median-unbiased and ML estimators of random variables are not necessarily the median-unbiased or ML estimators of sum of these variables (which applies to multi-nucleotide region calculations). This means that in general instead of summation of estimators, one has to recalculate the estimator for the whole sum. Luckily for the case of nucleotide diversity if a region has more SNPs than the individuals in the sample – the mean-unbiased estimator becomes median-unbiased (though not ML). Yet impossible values can appear, but they can be manually removed without damage to the median-unbiasedness.

We also developed MLE and median-unbiased methods for calculating Watterson θ. While standard Watterson estimator estimates the mean, and it does not depend on values of N and *μ* but only on their product, both MLE and median-unbiased method have such dependence. However, when *μ* <<1 this dependence is negligible and, therefore, we can substitute any large number for N to find θ. We also note that Watterson θ is a score with very high error which cannot be fully removed by increasing sample size. Still, the proposed MLE method is shown to reach smaller median error faster than the standard estimator and thus smaller datasets are required for it to reach accurate estimates. For the standard estimator the error is such that for 50% of cases and sample size 100 the value is over 5% less than the true value. The high error of Watterson θ was addressed by Futschik ^13^ and Fu ^14^. However, their methods are based on SFS and therefore heavily depend on the assumptions of the absence of selection and constant population size, and thus they become less accurate if these assumptions are broken. For the same reason, such type of estimator cannot be used for Tajima’s D. Moreover, for the BLE in Futschik and Gach 2008, we observe a much larger median error and median bias than for all other discussed θ estimators. In case of high recombination, mutation distribution turns into Poisson with standard Watterson estimator being MLE, and Fu’s estimator also coincides with the standard estimator because the error cannot be improved.

We also improve estimators by taking into account the uncertainty in the genotype calling through using Phred scores. Previously, Johnson and Slatkin 2006^9^ developed a Watterson θ estimator based on Phred scores. However, although the authors claimed to calculate an ML estimate, they actually calculated an unbiased (i.e the mean) estimate because of the assumed site independence. Furthermore, they calculated the probability that site is not polymorphic as 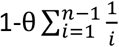 which is not quite correct as Wright-Fisher model assumes infinite number of sites^17^ and thus a 0 probability for a certain site to be polymorphic. Another work by Johnson and Slatkin 2008^10^ claims to produce unbiased estimators for θ (and π). However, they use average basecall error probability ε for all nucleotides in a region instead of individual error probabilities which reduces accuracy and causes bias. This is clearly seen from equation (2) in the paper. In ANGSD^1^ authors propose using ML or empirical Bayes estimates of SFS for calculations of population genetic statistics. The resulting estimate however is neither ML nor unbiased^7^.

The high error of Watterson θ and median-bias affect statistics that use it, such as Tajima’s D. Because θ and π are close values, the error for D is likely to achieve even larger error than the true value. Tajima’s D has negative median bias, meaning that most scores in neutral case are negative. Moreover, Tajima’s π is not a consistent estimator, meaning that the error would be large even for infinite sample. Finally, Tajima’s D depends on mutation rate which means that it can produce different results for populations with the same history and selection but different mutation rates. For these reasons, we propose another estimator D’ which does not suffer from this problem and can detect smaller deviations from neutral model.

We are looking into applying these methods to metagenome/pooled data. In such data each read may come from a different individual which affects likelihoods and statistics^20^. Moreover, Watterson θ and therefore Tajima’s D require us to know the sample size which is not known in metagenomes and is different for different species (and effectively even genome regions). In our future work we will be providing solutions to these questions as well as look closer into four allele ML and median-unbiased estimators for ***π***.

A tool implementing the discussed methods is available at: https://github.com/anulin/PiThetic

## References

1. Korneliussen, T. S., Albrechtsen, A. & Nielsen, R. ANGSD: Analysis of Next Generation Sequencing Data. BMC Bioinformatics 15, 356 (2014).

2. Poplin, R. et al. Scaling accurate genetic variant discovery to tens of thousands of samples. 201178 Preprint at 10.1101/201178 (2018).

3. Garrison, E. & Marth, G. Haplotype-based variant detection from short-read sequencing. Preprint at 10.48550/arXiv.1207.3907 (2012).

4. Korunes, K. L. & Samuk, K. pixy: Unbiased estimation of nucleotide diversity and divergence in the presence of missing data. Molecular Ecology Resources 21, 1359–1368 (2021).

5. Chang, C. C. et al. Second-generation PLINK: rising to the challenge of larger and richer datasets. GigaScience 4, s13742-015-0047–8 (2015).

6. Danecek, P. et al. The variant call format and VCFtools. Bioinformatics 27, 2156–2158 (2011).

7. Korneliussen, T. S., Moltke, I., Albrechtsen, A. & Nielsen, R. Calculation of Tajima’s D and other neutrality test statistics from low depth next-generation sequencing data. BMC Bioinformatics 14, 289 (2013).

8. Quality Scores for Next-Generation Sequencing.

9. Johnson, P. L. F. & Slatkin, M. Inference of population genetic parameters in metagenomics: A clean look at messy data. Genome Res. 16, 1320–1327 (2006).

10. Johnson, P. L. F. & Slatkin, M. Accounting for Bias from Sequencing Error in Population Genetic Estimates. Molecular Biology and Evolution 25, 199–206 (2008).

11. Nei, M. & Li, W. H. Mathematical model for studying genetic variation in terms of restriction endonucleases. Proc. Natl. Acad. Sci. U.S.A. 76, 5269–5273 (1979).

12. Yang, Z. & Bielawski, J. P. Statistical methods for detecting molecular adaptation. Trends Ecol Evol 15, 496–503 (2000).

13. Futschik, A. & Gach, F. On the inadmissibility of Watterson’s estimator. Theoretical Population Biology 73, 212–221 (2008).

14. Fu, Y. X. Estimating effective population size or mutation rate using the frequencies of mutations of various classes in a sample of DNA sequences. Genetics 138, 1375–1386 (1994).

15. Ferretti, L. & Ramos-Onsins, S. E. A generalized Watterson estimator for next-generation sequencing: From trios to autopolyploids. Theoretical Population Biology 100, 79–87 (2015).

16. DePristo, M. A. et al. A framework for variation discovery and genotyping using next-generation DNA sequencing data. Nat Genet 43, 491–498 (2011).

17. Watterson, G. A. On the number of segregating sites in genetical models without recombination. Theor Popul Biol 7, 256–276 (1975).

18. Tajima, F. Statistical Method for Testing the Neutral Mutation Hypothesis by DNA Polymorphism. Genetics 123, 585–595 (1989).

19. Tajima, F. Evolutionary relationship of DNA sequences in finite populations. Genetics 105, 437–460 (1983).

20. Kofler, R. et al. PoPoolation: A Toolbox for Population Genetic Analysis of Next Generation Sequencing Data from Pooled Individuals. PLOS ONE 6, e15925 (2011).

