## Supplementary Figures for "Maximum likelihood point estimates for improved population genetics statistics"

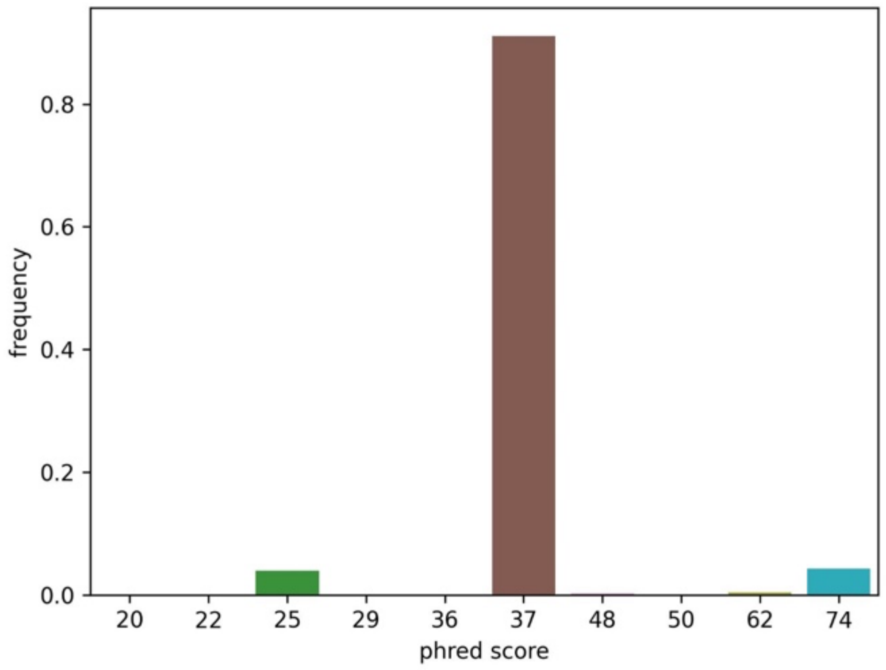


***Figure S1.*** Base quality score distribution.

#### Allele frequency estimation for region statistic estimators

For use in region statistic estimators for each allele pair frequency $\hat{p}$ is estimated as the root of the equation (15) where $p_i$ is the probability of observed read given reference allele, $p`_i$ is the probability of observed read given reference or alternative allele and $p``_i$ is the probability of observed read given alternative allele.

If the allele pair is {A,T} then for observed i$_{th}$ read A:

$$p_i = h_i;\ p`_i = \frac{2\mathrm{h_i} + 1}{3}; p``_i = \frac{1 - h_i}{3}$$

T:

$$p_i = \frac{1 - h_i}{3};\ p`_i = \frac{2\mathrm{h_i} + 1}{3}; p``_i = h_i$$

G and C:

$$p_i = \frac{1 - h_i}{3};\ p`_i = \frac{1 - h_i}{3}; p``_i = \frac{1 - h_i}{3}$$

Same applies to all other allele pairs.

To indicate value for $j_{th}$ genome we will use $p_{i,j}, p`_{i,j}$ and $p``_{i,j}$. Probability $p_{ab}$ of allele pair *ab* given the observed data is estimated as:

$$p_{ab} = \frac{P(observed|ab)}{\sum_{xy} P(observed|xy)} = \frac{\prod_j^N \left(p^2 \prod_i p_{i,j} + \frac{p*(1-p)}{2^{c-1}} \prod_i p`_{i,j} + (1-p)^2 \prod_i p``_{i,j} \,\middle|\, ab\right)}{\sum_{ij} \prod_j^N \left(p^2 \prod_i p_{i,j} + \frac{p*(1-p)}{2^{c-1}} \prod_i p`_{i,j} + (1-p)^2 \prod_i p``_{i,j} \,\middle|\, xy\right)}$$

*N* is the number of individuals.

#### Watterson θ

For diploid organism the number of genomes n=2*N*.

Mutation number for θ is estimated as mutation probability in each position:

$$P_{mut} = 1 - \prod_k p_{11,k} - \prod_k p_{00,k}$$

Where $p_{11,k}$ and $p_{00,k}$ are probabilities that $k_{th}$ individual is homozygous for reference allele and for alternative allele respectively, given the *ab* allele pair.

$$p_{11,k} = \left(\frac{p^2 \prod_i p_{i,k}}{p^2 \prod_i p_{i,k} + \frac{p(1-p)}{2^{c-1}} \prod_i p`_{i,k} + (1-p)^2 \prod_i p``_{i,k}} \,\middle|\, ab\right)$$

$$p_{00,k} = \left(\frac{(1-p)^2 \prod_i p``_{i,k}}{p^2 \prod_i p_{i,k} + \frac{p(1-p)}{2^{c-1}} \prod_i p`_{i,k} + (1-p)^2 \prod_i p``_{i,k}} \,\middle|\, ab\right)$$

For each position $m$ average $\hat{\theta}_m$ is calculated over possible allele pairs:

$$\hat{\theta}_m = \sum_{ab} p_{ab} \left(\frac{P_{mut}}{H_j} \,\middle|\, ab\right)$$

Where $H_j$ is a harmonic number.

Variance of $\hat{\theta}_m$ is calculated as:

$$Var(\hat{\theta}_m) = \sum_{ab} p_{ab} \left(\frac{P_{mut}}{H_j^2} \,\middle|\, ab\right) - \hat{\theta}_m^2$$

Then total $\hat{\theta}$ is:

$$\hat{\theta} = \frac{\sum_m \frac{\hat{\theta}_m}{Var(\hat{\theta}_m)}}{\sum_m \frac{1}{Var(\hat{\theta}_m)}} * L$$

Where $L$ is the window size.

**Nucleotide diversity for region**

For regions PiThetic uses fast π estimation (from formula 16). For a given allele pair *ab* nucleotide diversity estimator is:

$$\hat{\pi}(\hat{p}) = \left(\sum_k \frac{X(\hat{p})_k}{N} - \left({\sum_k X(\hat{p})_k}^2 - \sum_k X(\hat{p})_k^2 + 2 * \sum_k p_{11,k}(\hat{p})\right)/N(2N-1)\right)$$

Here $\hat{p}$ is allele frequency estimate given the *ab* allele pair (from formula 15).

$$\widehat{Var}\,\pi(p) = \sigma_\pi^2(p) = E[\hat{\pi}^2|p] - \hat{\pi}(p)^2$$

$X(p)_k$ is expected allele count of $k_{th}$ individual given the allele frequency p.

$$X(p)_k = 2 * p_{11,k}(p) + p_{10,k}(p)$$

Probability to observe an exact set of reads from an individual given the allele frequency *p* is:

$$P(\{p_i\}, p) = p^2 \prod_i p_i + \frac{p(1-p)}{2^{c-1}} \prod_i p`_i + (1-p)^2 \prod_i p``_i$$

Here $\{p_i\}$ are the observed nucleotides (numbers of reads and phred scores are fixed). The expected value of $\hat{\pi}^2$ is:

$$E\hat{\pi}^2 = \sum_{\{p_i\}} P(p, \{p_i\}) * \hat{\pi}^2$$

This equation consists of terms $X_k(p)X_j(p), X_k p_{11,j}$ , $k! = j. X_k,\ X_j$ are independent so:

$$E[X_k X_j] = EX_k EX_j = 4p^2,$$

$X_k(p)p_{11,j}(p), j! = k;\ E[X_k p_{11,j}] = EX_k Ep_{11,j} = 2p^3$

terms $X_k^2$

$$EX_k^2 = \sum_{\{p_i\}} P(\{p_i\}, p) * X_k^2 = \sum_{\{p_i\}} P(\{p_i\}, p) \left(\frac{2 * p^2 \prod_i p_i + \frac{p*(1-p)}{2^{c-1}} \prod_i p`_i}{p^2 \prod_i p_i + \frac{p*(1-p)}{2^{c-1}} \prod_i p`_i + (1-p)^2 \prod_i p``_i}\right)^2$$

$$= \sum_{\{p_i\}} P(\{p_i\}, p) \left(\frac{p^2 \prod_i p_i - (1-p)^2 \prod_i p``_i}{p^2 \prod_i p_i + \frac{p*(1-p)}{2^{c-1}} \prod_i p`_i + (1-p)^2 \prod_i p``_i} + 1\right)^2$$

$$= 4p - 1 + \sum_{\{p_i\}} \frac{(p^2 \prod_i p_i - (1-p)^2 \prod_i p``_i)^2}{p^2 \prod_i p_i + \frac{p*(1-p)}{2^{c-1}} \prod_i p`_i + (1-p)^2 \prod_i p``_i}$$

And also terms $X_k p_{11,k}$:

$$EX_k p_{11,k} = \sum_{\{p_i\}} P(\{p_i\}, p) X_k p_{11,k}$$

$$= \sum_{\{p_i\}} P(\{p_i\}, p) \left( \frac{p^2 \prod_i p_i - (1-p)^2 \prod_i p``_i}{p^2 \prod_i p_i + \frac{p*(1-p)}{2^{c-1}} \prod_i p`_i + (1-p)^2 \prod_i p``_i} + 1 \right) p_{11,k} =$$

$$= p^2 + \sum_{\{p_i\}} \frac{p^4 \prod_i p_i^2 - (1-p)^2 p^2 \prod_i p``_i p_i}{p^2 \prod_i p_i + \frac{p*(1-p)}{2^{c-1}} \prod_i p`_i + (1-p)^2 \prod_i p``_i}$$

Now because the size of $\{p_i\}$ is $4^{coverage}$ and there is no obvious shortcut for this formula, it has to be computed directly which gets exponentially hard with growing coverage. To overcome this, we only calculate values for cases when there are 0 or 1 erroneous (and also 0 or 1 correct because the formula is symmetric) reads. When there is no error the term is

$$X_{0,m}(p) = \frac{\left(p^2 \prod_i p_i - (1-p)^2 \prod_i p``_i\right)^2}{p^2 \prod_i p_i + \frac{p*(1-p)}{2^{c-1}} \prod_i p`_i + (1-p)^2 \prod_i p``_i}$$

When there is one error the terms are:

$$X_{1,m}(p) = \sum_j \frac{\left(\frac{1-p_i}{3p_i} p^2 \prod_i p_i - \frac{3p_i}{1-p_i}(1-p)^2 \prod_i p``_i\right)^2}{\frac{1-p_i}{3p_i} p^2 \prod_i p_i + \frac{p*(1-p)}{2^{c-1}} \prod_i p`_i + \frac{3p_i}{1-p_i}(1-p)^2 \prod_i p``_i}$$

$$+ 2\sum_j \frac{\left(\frac{1-p_i}{3p_i} p^2 \prod_i p_i - \frac{3p_i}{1-p_i}(1-p)^2 \prod_i p``_i\right)^2}{\frac{1-p_i}{3p_i} p^2 \prod_i p_i + \frac{(1-p_i)p*(1-p)}{(2p_i+1)2^{c-2}} \prod_i p`_i + \frac{3p_i}{1-p_i}(1-p)^2 \prod_i p``_i}$$

Therefore, the approximate $EX_k^2$:

$$EX_k^2 \approx 4p - 1 + X_{0,m}(p) + X_{1,m}(p) + X_{0,m}(1-p) + X_{1,m}(1-p)$$

$EX_k p_{11,k}$ and $Ep_{11,k}^2$ are calculated with the same logic.

$$E\hat{\pi}(p)^2 = E\left(\sum_k \frac{X(p)_k}{N} - \frac{\left(\sum_{k!=j} X(p)_k X(p)_j + 2*\sum_k p_{11,k}(p)\right)}{N(2N-1)}\right)^2 = a + b + c$$

$$a = \sum_k \frac{EX_k^2}{N^2} + E\sum_{k!=j} \frac{X_k X_j}{N^2} - 2E\sum_{k!=j} \frac{X_k^2 X_j}{N^2(2N-1)} - 2E\sum_{k!=j!=m} \frac{X_k X_j X_m}{N^2(2N-1)} - \frac{4E\sum_k X_k p_{11,k}}{N^2(2N-1)}$$

$$- 4\sum_{k!=j} \frac{X_k p_{11,j}}{N^2(2N-1)}$$

$$= \sum_k \left(\frac{EX_k^2}{N^2} - 4\frac{EX_k^2 p(N-1)}{N^2(2N-1)} - \frac{4EX_k p_{11,k}}{N^2(2N-1)}\right) + \frac{N-1}{N}4p^2$$

$$- \frac{16p^3(N-1)(N-2)}{N(2N-1)} - 8\frac{p^3(N-1)}{N(2N-1)}$$

$$a = \sum_k \frac{EX_k^2\big(2N-1-4p(N-1)\big) - 4EX_k p_{11,k}}{N^2(2N-1)} + \frac{N-1}{N}4p^2 - \frac{8p^3(N-1)(2N-3)}{N(2N-1)}$$

$$b = \frac{E\left(\sum_{k!=j!=m!=q} X_k X_j X_m X_q + \sum_{k!=j!=m}\left(X_k^2 X_j X_m + 4X_j X_m p_{11,k}\right) + \sum_{k!=j}(X_k^2 X_j^2 + 4X_j X_k p_{11,k})\right)}{N^2(2N-1)^2}$$

$$b = \frac{16p^4(N-1)(N-2)^2}{N(2N-1)^2} + \frac{\sum_k\left(EX_k^2 4p^2(N-1)(N-2) + 8p(N-1)EX_k p_{11,k}\right) + \sum_{k!=j} EX_k^2 EX_j^2}{N^2(2N-1)^2}$$

$$c = \frac{4E\left(\sum_k p_{11,k}^2 + \sum_{k!=j} p_{11,k} p_{11,j}\right)}{N^2(2N-1)^2} = \frac{4\sum_k Ep_{11,k}^2}{N^2(2N-1)^2} + \frac{4(N-1)p^4}{N(2N-1)^2}$$

Then given that allele pair is not known we get:

$$\hat{\pi} = \sum_{ab} p_{ab}(\hat{\pi}|ab) = \sum_{ab} p_{ab}\hat{\pi}_{ab}$$

$$E\hat{\pi}^2 = \sum_{ab} p_{ab}\hat{\pi}_{ab}^2$$

$$\widehat{Var(\pi)} = E\hat{\pi}^2 - \hat{\pi}^2 = \sigma_\pi^2(\hat{\boldsymbol{p}})$$

Here $\hat{\boldsymbol{p}}$ is a vector of $\hat{p}$ estimates for all allele pairs ab. For locus j $\sigma_\pi^2$=$\sigma_{\pi j}^2$. Then total $\hat{\pi}$ for a region is calculated as:

$$\hat{\pi}_{total} = \sum_j^l \left( \frac{\frac{\hat{\pi}_j(\hat{\boldsymbol{p}}_{\boldsymbol{j}})}{\sigma_{\pi j}^2(\hat{\boldsymbol{p}}_{\boldsymbol{j}})}}{\sum_i^N \frac{1}{\sigma_{\pi i}^2(\hat{\boldsymbol{p}}_{\boldsymbol{j}})}} \right)$$

Because this can be computationally long to estimate variances for each frequency for each combination of reads ($O(l^2)$ where $l$ is the length of window) we also provide a fast less accurate option in our tool:

$$\hat{\pi}_{fast} = \sum_j^l \hat{\pi}_j(\hat{\boldsymbol{p}}_{\boldsymbol{j}})$$

## 5. D’

$$D' = \frac{\frac{\hat{\pi} l}{K} - \frac{1}{\sum_{i=1}^{n-1}\frac{1}{i}}}{\sqrt{\widehat{Var\left(\frac{\pi l}{K}\right)}}}$$

Where K is the mutation number. Previously we explained how we estimate $\hat{\pi}$. $Var\left(\frac{\pi l}{K}\right)$ is estimated in the following way:

From [1]:

$$Var(\pi l) = E(\pi l)^2 - \theta^2 = \frac{n+1}{3(n-1)}\theta + \frac{2(n^2+n+3)}{9n(n-1)}\theta^2$$

From coalescent theory:

$$E(\pi l)^2 = E[K(K-1)]E_{i!=j}\pi_i\pi_j + EKE\pi_i^2$$

For $\frac{\pi l}{K}$ K is fixed, so:

$$E\left(\frac{\pi l}{K}\right)^2 = \frac{(K-1)}{K}E\pi_i\pi_j + \frac{1}{K}E\pi_i^2$$

$$E\pi_i^2 = \frac{n+1}{3(n-1)}$$

$$E\pi_i\pi_j = \frac{Var(\pi l) + \theta^2 - \frac{n+1}{3(n-1)}\theta}{E[K(K-1)]} = \frac{11\text{n}^2 - 7\text{n} + 6}{9n(n-1)E[K(K-1)]}\theta^2$$

From[2]:

$$E[K(K-1)] = \theta^2\left(\sum_i^{n-1}\frac{1}{i^2} + \left(\sum_i^{n-1}\frac{1}{i}\right)^2\right)$$

$$E\pi_i\pi_j = \frac{11\text{n}^2 - 7\text{n} + 6}{9n(n-1)\left(\sum_i^{n-1}\frac{1}{i^2} + \left(\sum_i^{n-1}\frac{1}{i}\right)^2\right)}$$

$$\widehat{Var\left(\frac{\pi l}{K}\right)} = \frac{(K-1)}{K}\frac{11\text{n}^2 - 7\text{n} + 6}{9n(n-1)\left(\sum_i^{n-1}\frac{1}{i^2} + \left(\sum_i^{n-1}\frac{1}{i}\right)^2\right)} + \frac{1}{K}\frac{n+1}{3(n-1)} - \frac{1}{\left(\sum_i^{n-1}\frac{1}{i}\right)^2}$$

Variance of mutation number in single position is calculated as:

$$Var(P_{mut}) = \sum_{ab} p_{ab}(P_{mut}\ |ab) - P_{mut}^2$$

Then K is:

$$K = \frac{\sum_m \frac{P_{mut}}{Var(P_{mut})}}{\sum_m \frac{1}{Var(P_{mut})}} * L$$

$$K = \sum_{ab} p_{ab}(P_{mut}\ |ab)$$

### Simulations with variable coverage

To simulate data, we sampled a set of 60 Phred scores from real-life sequencing data (distribution shown in **Supplementary Fig S1**), such that there were lower scores present in the sample. SNPs were assumed to be biallelic. Random values from binomial distribution with a probability *p* were sampled to get true sample allele frequencies. Phred scores were sampled randomly without replacement, and the erroneous calls were sampled from a Poisson-Binomial distribution, with the probabilities defined with the sampled Phred scores as parameters, and applied to each individual to simulate the observed reads. Diploid samples from a population of 15 individuals with sequencing coverage drawn from Poisson distribution (λ=4) were generated. Data generation was done 100,000 times for ten different values of *p* in an interval [0,0.5] with increment of 0.0(5). Likelihoods were calculated for the data using the Phred scores. Medians and means of standard (SNP call) allele frequency, MLE for frequency (13) were compared and ML frequency estimates are more accurate than the standard estimates(**Fig S2,3)**. Similar comparison but for errors was done for $\pi$ from formula (14), $\pi$ calculated from MLE frequency and the standard (naïve) $\pi$ (**Fig S4-6)**. In terms of median, ML $\pi$ estimates (14) are generally the most accurate. In all cases naïve $\pi$ has error several times bigger than MLE $\pi$ when $\pi < 0.2$ which is important because it is rare when $\pi$ exceeds 0.02.

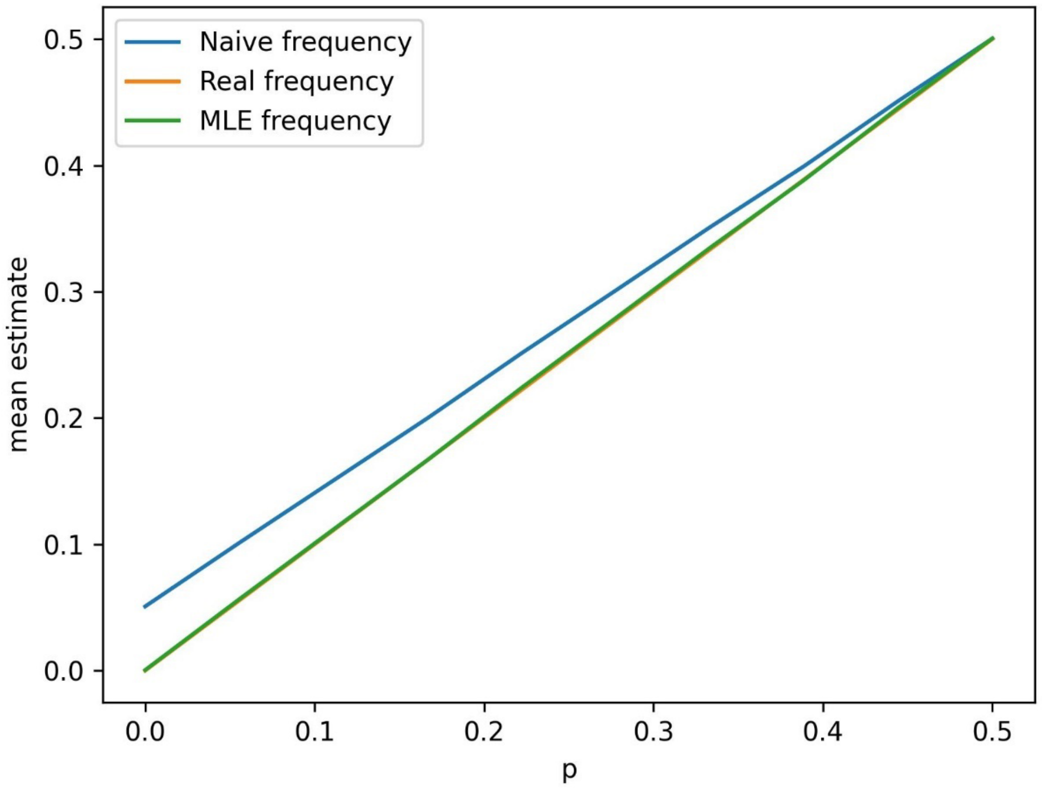


***Figure S2.*** Mean frequency estimators for simulations with random coverage.

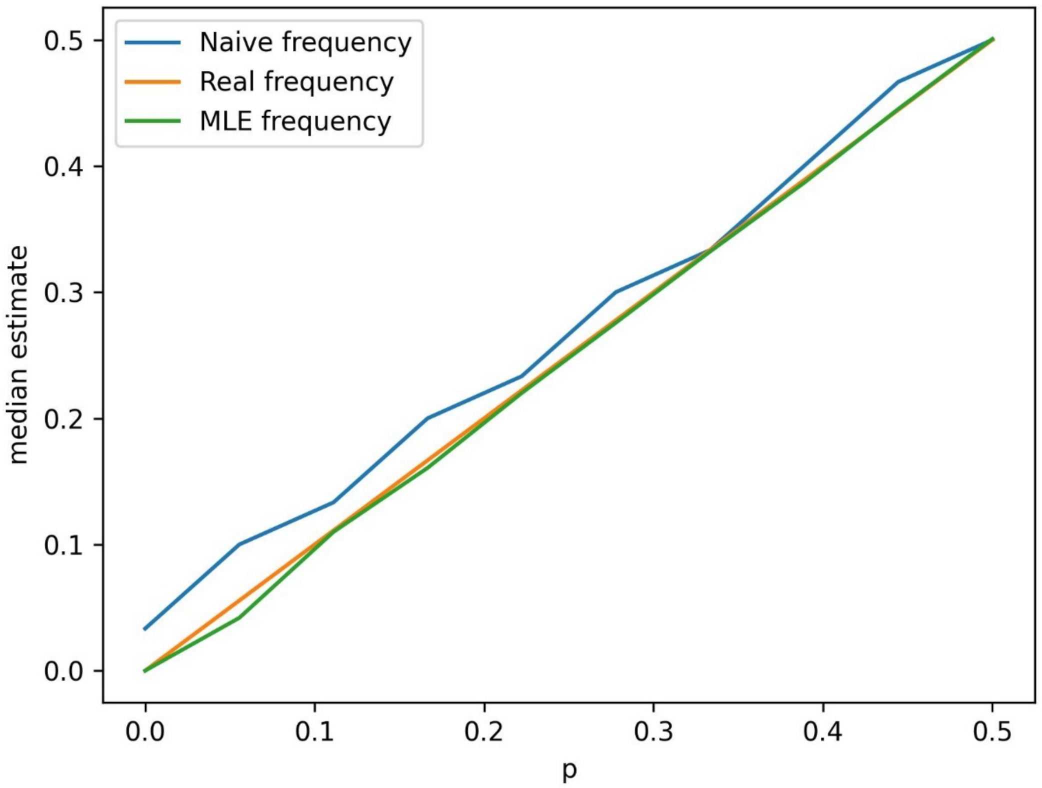


***Figure S3.*** Median frequency estimators for simulations with random coverage.

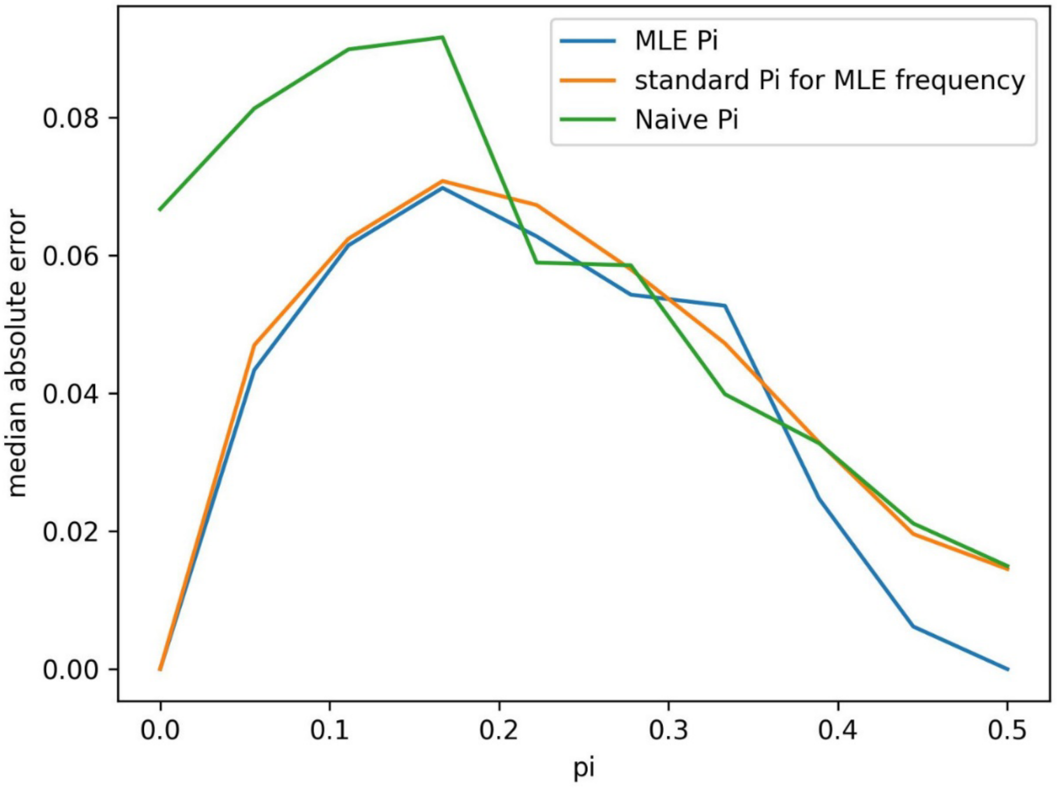

***Figure S4.*** π estimator median absolute errors for simulations with random coverage.

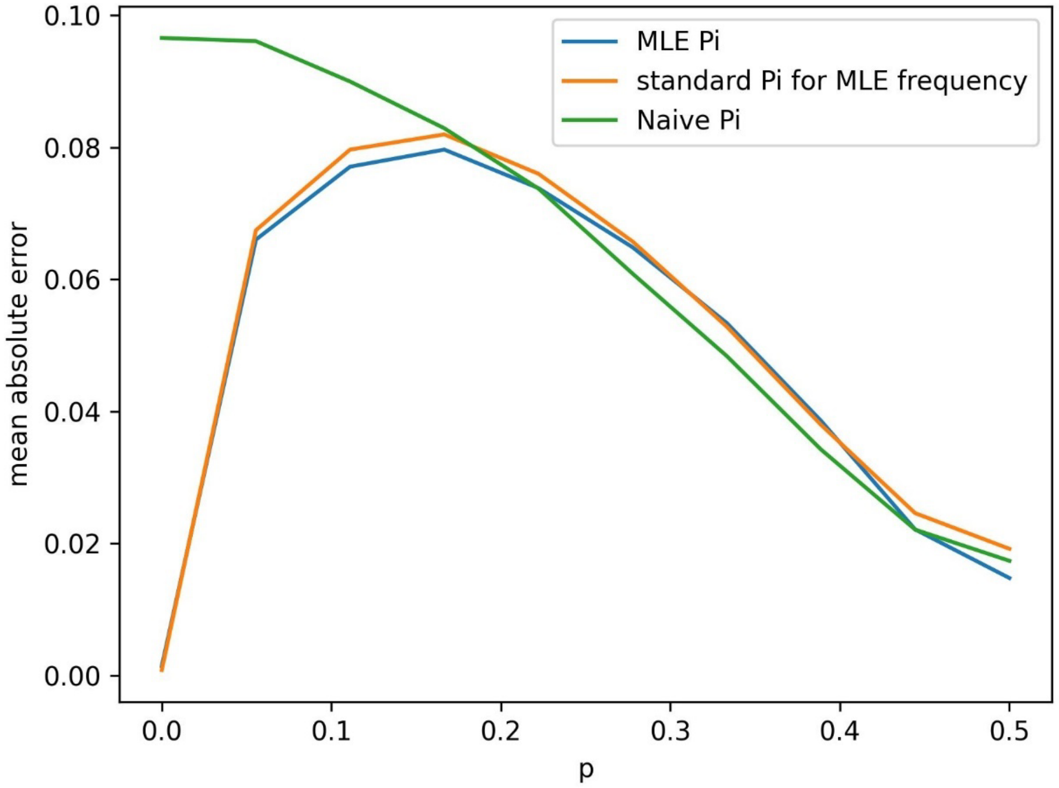


***Figure S5.*** π estimator mean absolute errors for simulations with random coverage.

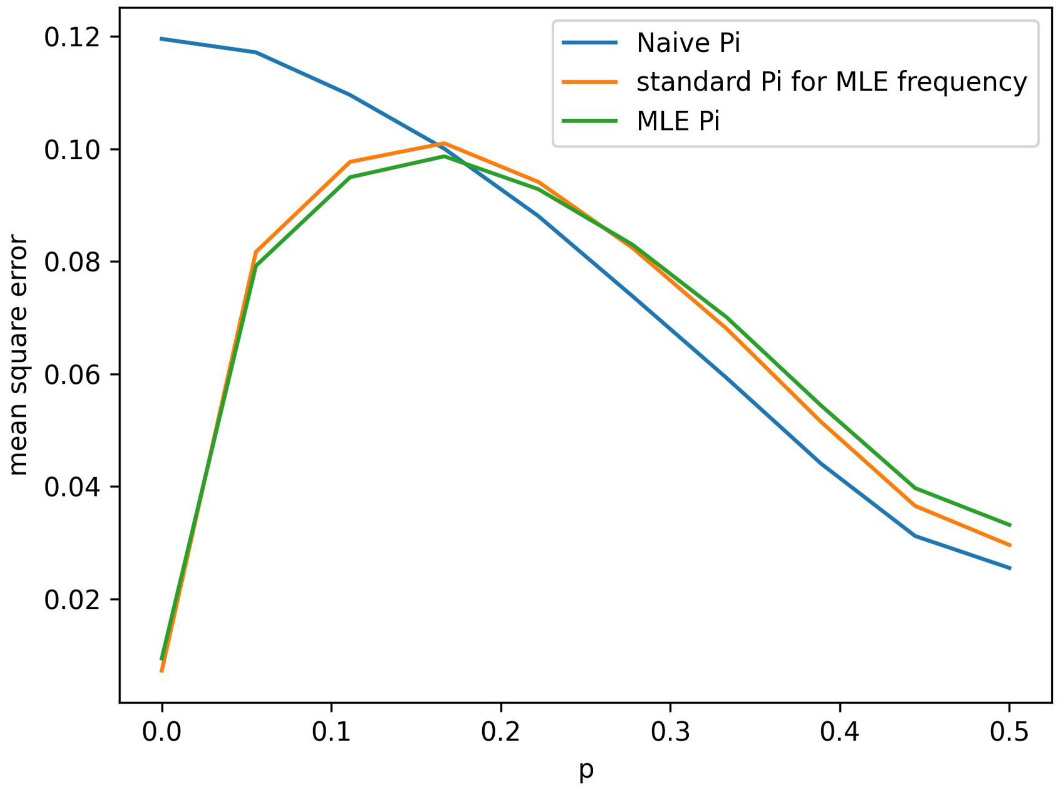


***Figure S6.*** π estimator standard errors for simulations with random coverage.
